# Spatiotemporal biogenesis of thylakoid membranes in the green alga *Chlamydomonas reinhardtii*

**DOI:** 10.64898/2026.06.18.733168

**Authors:** Xingwu Ge, Zhen Hou, Man Qi, Paula Muñoz, Krzysztof Pawlak, Gregory F. Dykes, Yingyue Zhang, Joscelyn Sarsby, Jianguo Zhang, Long Chen, Jessica Heebner, Maud Dumoux, Giles N. Johnson, Peter J. Nixon, Peijun Zhang, Lu-Ning Liu

## Abstract

Thylakoid membranes are essential for oxygenic photosynthesis, yet the mechanisms underlying their spatial and temporal biogenesis remain poorly understood. Here, using a light-induced membrane regeneration system in *Chlamydomonas reinhardtii*, we generate a time-resolved map of thylakoid formation and photosynthetic complex assembly at unprecedented spatial and temporal resolution by integrating super-resolution fluorescence microscopy, cryo-electron tomography, proteomics, and spectroscopy. We show that thylakoid biogenesis is globally distributed and multipolar, with new membrane formation occurring at multiple sites across the chloroplast, including regions adjacent to the inner envelope membranes in both basal and lobe regions. We identify F-ATP synthase storage membranes and map the redistribution of translating ribosomes from T-zone enrichment to a chloroplast-wide distribution at early stages of thylakoid formation, consistent with decentralized synthesis of photosynthetic complexes. We further delineate the hierarchical assembly of photosystem supercomplexes, starting with formation of reaction centers. Together, these findings refine the spatial organization of photosynthetic membrane biogenesis and establish a unified spatiotemporal framework linking thylakoid formation, translational activation, and functional maturation, providing a mechanistic basis for plastid engineering to enhance photosynthetic efficiency.

## Introduction

Life on Earth depends on oxygenic photosynthesis, which is a vital biological process that harnesses solar radiation to generate chemical energy while producing oxygen. The thylakoid membrane, which is intricately organized within chloroplasts and cyanobacteria, serves as a specialized platform for the light-dependent reactions of oxygenic photosynthesis ^1–3^. In the thylakoid membrane, the central photosynthetic complexes, including photosystem II (PSII), photosystem I (PSI), cytochrome (Cyt) *b_6_f* complex, and ATP synthase (F-ATPase), mediate light-driven electron transport to form NADPH and the transmembrane proton gradient essential for ATP synthesis ^4,5^. These energy-rich products power cell metabolism, such as the Calvin-Benson cycle, thus linking sunlight capture to the biosynthesis of photoautotrophic cells.

In eukaryotic algae and plants, thylakoid membranes are arranged in the specialized organelle called the chloroplast ^5–8^. The thylakoid architecture in vascular plants exhibits a high degree of structural complexity, characterized by juxtaposition of stacked grana and unstacked stroma lamellae. In contrast, green algae such as *Chlamydomonas reinhardtii* possess a less compartmentalized yet distinctly ordered thylakoid system closely integrated with the pyrenoid, a specialized microcompartment vital for carbon fixation ^9^. This structural complexity requires the sophisticated regulation of thylakoid membrane biogenesis and photosynthetic complex assembly to maintain photosynthetic efficiency and enables adaptive responses to environmental fluctuations ^10^. Despite their physiological importance, the fundamental molecular and cellular processes governing the biogenesis and regulation of thylakoid membranes remain elusive.

One long-standing question is the biogenic site of thylakoid membranes. Over the past decades, *C. reinhardtii* has been a model system for studying thylakoid biogenesis, given that it possesses a single chloroplast that occupies 40% of the cell volume and features typical morphology ^11,12^. The current model proposes that *de novo* photosystem biogenesis and chlorophyll (Chl) biosynthesis in *C. reinhardtii* can be concentrated in a specific “translation zone” (T-zone) in the outer perimeter of the pyrenoid within the chloroplast ^13–17^, although earlier electron microscopy studies also documented envelope-associated thylakoid formation during greening ^18,19^. The assembled photosynthetic supercomplexes then move laterally along contiguous thylakoid membrane layers to other regions of the chloroplast ^16,20^. However, the fine-scale spatial and temporal coordination of thylakoid initiation, expansion, and complex assembly during greening remains to be fully elucidated.

In this study, we established a system enabling extensive depletion of thylakoid membranes in *C. reinhardtii* and reactivation of thylakoid biogenesis upon light exposure. By integrating thin-section transmission electron microscopy (TEM), super-resolution fluorescence microscopy, in-cell cryo-electron tomography (cryo-ET) coupled with cryo-focused ion beam and scanning electron microscopy (cryo-FIB/SEM) volume imaging, proteomics, and spectroscopy, we performed a systematic, time-resolved analysis of *in vivo* thylakoid membrane biogenesis and the hierarchical assembly of photosynthetic complexes during the greening process. Our results reveal that thylakoid membrane biogenesis in *C. reinhardtii* proceeds as a globally distributed and multipolar process, rather than being confined to the previously hypothesized T-zone. These findings provide quantitative, molecular evidence for the spatiotemporal orchestration and regulation of thylakoid membrane biogenesis in algae and plants, thereby advancing the conceptual framework for understanding chloroplast development.

## Results

### Generation of a *chlL*-deletion mutant for studying the greening process of *Chlamydomonas*

Previous studies on thylakoid membrane biogenesis in *Chlamydomonas* were performed by examining the greening of a *yellow-in-the-dark-1* (*y-1*) mutant ^14,18,19,21,22^. The *y-1* mutant blocks light-independent Chl biosynthesis due to the absence of the dark-operative protochlorophyllide oxidoreductase (DPOR) required for Chl formation in the dark (Fig. S1a). However, the exact nature of the nuclear mutation in *y-1* remains unclear ^23^, and exhibits genetic instability ^24^. Apart from *y-1*, disruption of other three chloroplast genes (*chlL*, *chlN*, and *chlB*) encoding DPOR produced mutants displaying a yellow-in-the-dark phenotype ^23,25–28^, offering alternative solutions for exploring thylakoid membrane biogenesis in *Chlamydomonas*.

We generated a Δ*chlL* strain by deleting the *chlL* gene from the plastid genome (Figs. S1b, S1c). To initiate the greening process, the Δ*chlL* strain was grown heterotrophically in the dark in Tris-acetate-phosphate (TAP) medium with acetate as the carbon source for more than four generations prior to light treatment (Fig. 1a). In dark-grown cells (0 h), the Chl concentration in the Δ*chlL* cells was substantially lower than in the wild type (WT) cells, resulting in a yellowish colour in the dark (Figs. 1a-c, Fig. S2). Notably, the contents of Chl *a* and Chl *b* in Δ*chlL* grown in the dark were below the detectable levels via high-performance liquid chromatography (HPLC) (Fig. 1c). Analysis over a 24 h period showed that Chl synthesis in cells was rapidly activated by the light illumination (from 0.61 ± 0.38 to 17.78 ± 2.15 μg/mL/OD_750_, Fig. S2c), in agreement with the greening process described in previous studies ^18,29^. Both Chl *a* and Chl *b* were detectable within 4 h, along with an increase in the content of various Chl-like molecules, presumably Chl precursors (Fig. 1c), indicating active alterations in Chl biosynthesis and modification during the early greening process. Measurements of PSII maximum quantum efficiency (Fv/Fm) and oxygen evolution, as well as near-infrared absorbance changes reporting on P700 turnover, indicated that both PSII and PSI activities were largely diminished in the dark and at early stages and were gradually restored during greening (Figs. 1d, 1e, Fig. S2d). After 24 h of illumination, the Chl content and photosystem activities in Δ*chlL* were substantially recovered, approaching the levels in the WT (Figs. 1c-1e, Figs. S2a, S2c).

**Fig. 1.**
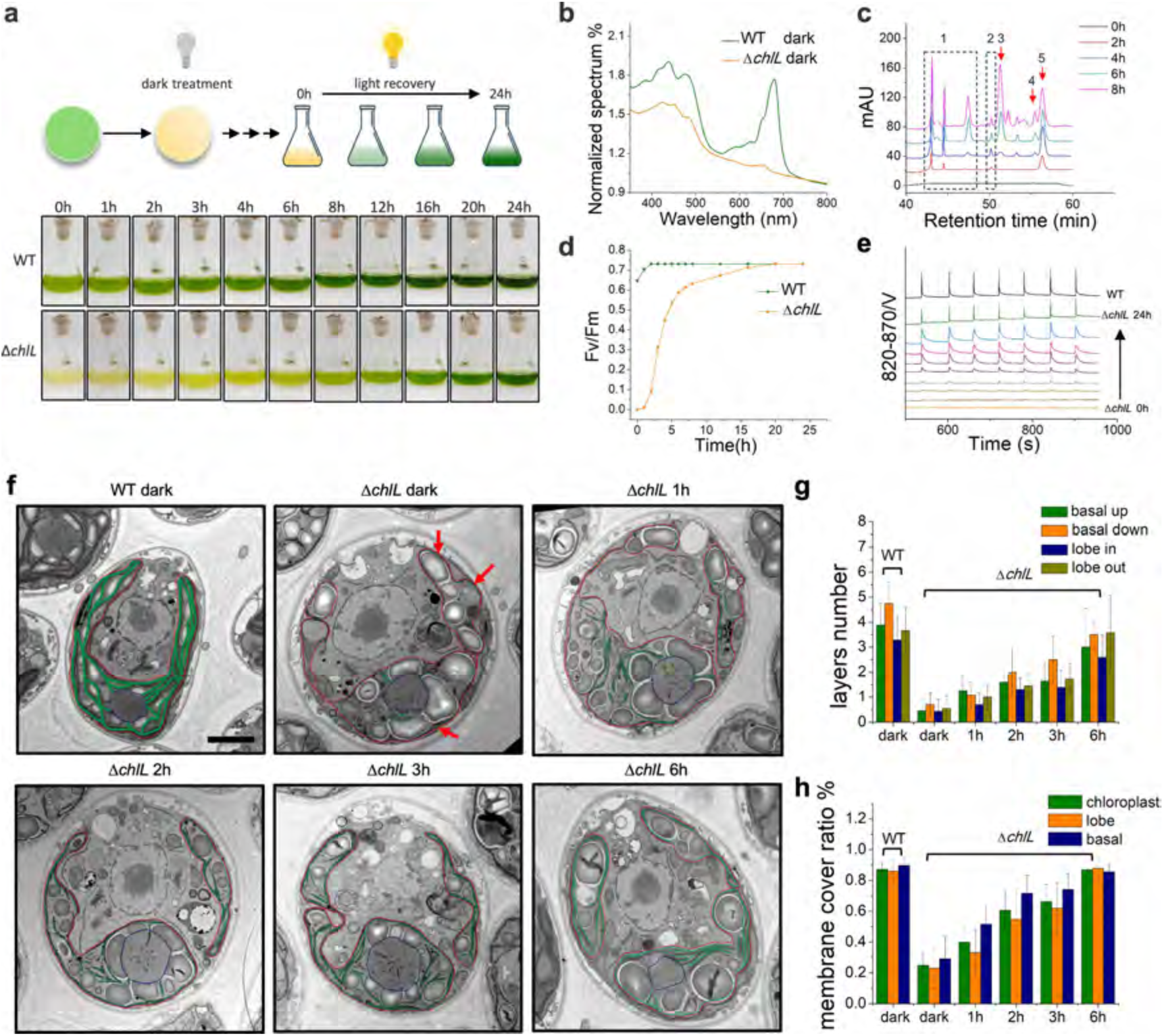
Characterization of thylakoid biogenesis from a thylakoid-less state during greening. **a**, Schematic of the greening procedure. *C. reinhardtii* cells were maintained on plates for over four generations until Chl levels fell below the detection threshold. ‘Yellow’ cultures were then recovered under low light (20 µmol photons m⁻² s⁻¹). **b**, Whole-cell absorbance spectra of 1 ml cultures (3 biologically independent replicates), normalized at 750 nm. **c**, HPLC analysis of pigments using C-18 reverse column, peaks 1∼5 were identified to be chlorophyll *a*-like, chlorophyll *b*-like, chlorophyll *a*, chlorophyll *b*, and chlorophyll *a*-like, respectively. **d**, Fv/Fm during the greening process. Data represent means ± SD; *n* = 3 biologically independent preparations. **e**, P700 slow kinetics measured by Dual KLAS NIR system. The absorbance changes in 820-870nm produced by each excitation represent the re-reduction activity of P700, Orange to green represent Δ*chlL* cells after light treatment for: 0 h (dark), 1 h, 2 h, 4 h, 6 h, 8 h, 12 h, 16 h, 24 h, the black line is measured from the light grown CC-1690. Offset was given to show each curve; the representative curve with seven excitations replications was from three replicates. **f**, Thin-section transmission electron microscopy (TEM) images of *C. reinhardtii* WT and the Δ*chlL* mutant. Cells were treated as described in (a) and immediately fixed at each time point. The chloroplast envelope is outlined in red; straight, smooth thylakoid membranes are marked in green. Red arrows indicate residual membranes in dark-treated Δ*chlL* cells. Scale bar = 2 µm. **g**, the number of thylakoid membrane layers near the chloroplast envelope determined in EM images. Values are means ± SD, *n* > 20. **h**, Ratio of the envelope length that with/without parallel thylakoid membranes determined in EM images. Values are means ± SD, *n* > 20.

### Development of thylakoid membranes during the greening process

We then sought to study *de novo* thylakoid membrane biogenesis using the Δ*chlL* strain. The Δ*chlL* cells were collected during the 0-6 h light treatment and imaged by thin-section TEM. Compared to the WT, the Δ*chlL* mutant displayed a remarkably low thylakoid membrane content in the dark (0 h) (Fig. 1f, Fig. S3a), aligning with the reduced pigment content detected in Δ*chlL* (Figs. 1a-1c, Figs. S2a, S2c). Typical stacked thylakoid membranes were absent in the Δ*chlL* chloroplast, with only residual and fragmented membrane structures remaining near the chloroplast inner envelope membrane (IEM), the stroma adjacent to the pyrenoid, and the stromal regions of the chloroplast lobes that are extended regions surrounding the nucleus and cytoplasm ^13,30^. In this study, we define these light-responsive precursor structures as nascent thylakoids (NTs), which are distinguished from mature continuous thylakoids by their size, fragmentation, and irregular curvature. Despite the depletion of thylakoid membrane content and reorganization of its architecture, the cells retained their characteristic internal structures ^9,11,30^, including a single cup-shaped chloroplast, the pyrenoid, and pyrenoid tubules (Fig. S3b).

In Δ*chlL* at 0 h, only ∼24% of the envelope regions displayed a single thylakoid membrane structure parallel to the chloroplast envelope, in contrast to ∼87% in the WT (Fig. 1g). To evaluate the membrane content, we further calculated the proportion of envelope regions possessing parallel thylakoid membranes. The IEM surface with parallel thylakoid membranes immediately increased in response to light exposure. During the early greening process, this proportion increased progressively; by 6 h, it reached 87%, which was comparable to that in the WT (Fig. 1h). Moreover, our results indicated that light also triggered a progressive increase in thylakoid membrane layers adjacent to the chloroplast IEM in both the basal and lobe regions (Fig. S3c), suggesting that thylakoid membrane development may occur at several distinct sites within the chloroplast.

### The biogenic sites of thylakoid membranes are globally distributed within the chloroplast

To probe the biogenic sites of thylakoid membranes within the chloroplast, we employed cryo-FIB/SEM to image the Δ*chlL C. reinhardtii* cells at 0 h, 1 h and 24 h in comparison with the WT in the dark (Table S1). In the reconstructed 3D cell volume (Fig. S4), drastic differences in the morphology and number of thylakoids were observed among the four samples. Consistent with the thin-section TEM data, the WT cells in dark preserved crowded stacks of continuous and appressed thylakoid membranes, whereas the Δ*chlL* mutant cells in the early stage (0-1 h) of biogenesis contained remarkably fewer, smaller, and fragmented NTs (Fig. 2a-b, Fig. S4, Movies S1-S4). After 24 h of light treatment, thylakoid membranes largely recovered and showed no differences in the overall morphology compared to the WT. In particular, the number of NTs increased more than three-fold in chloroplasts after 1 h of illumination (Fig. 2c, Fig. S4). In the 3D cell volume of Δ*chlL* cells, NTs were found across the entire chloroplast without specific localization.

**Fig. 2.**
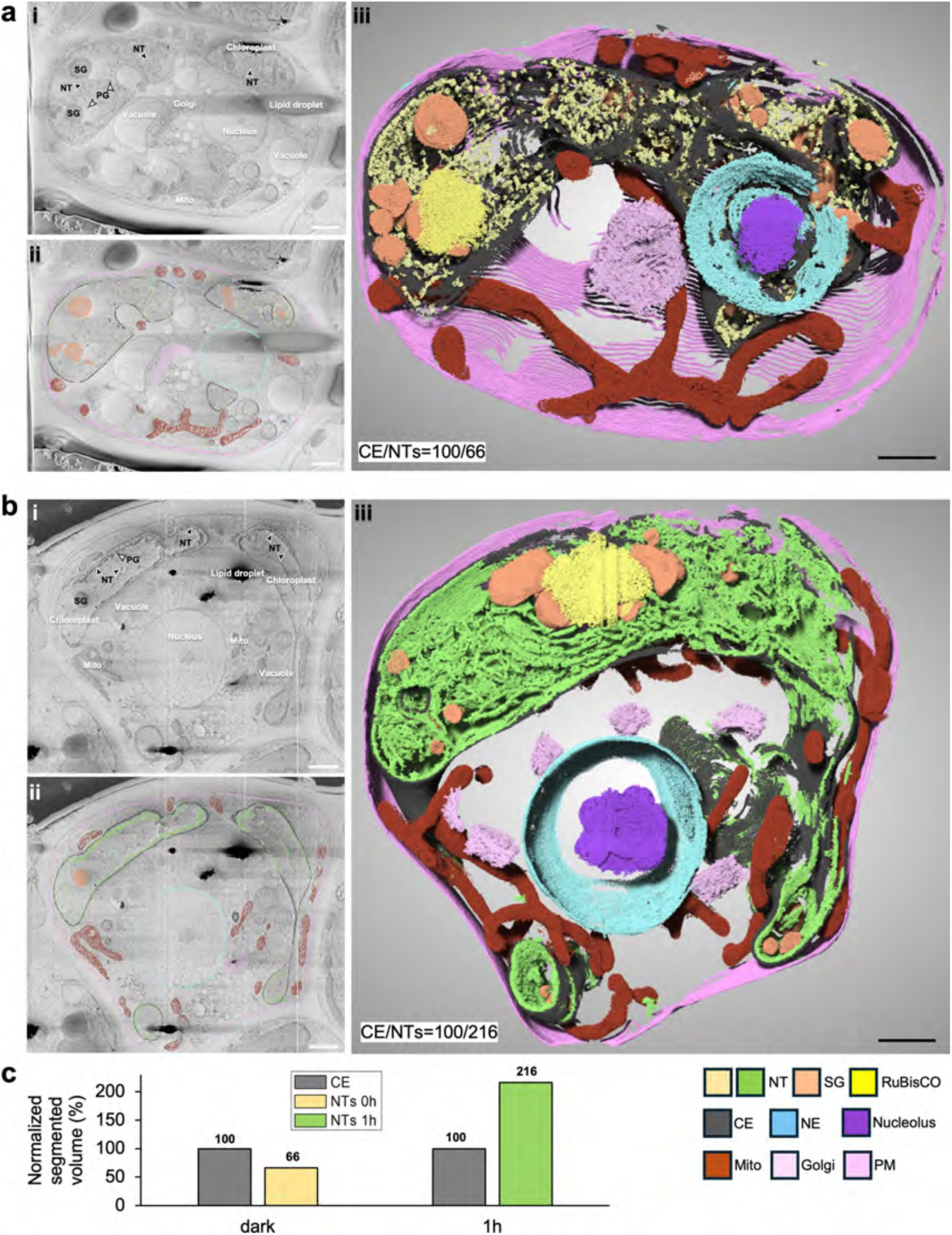
Nascent thylakoids (NTs) revealed in whole native chloroplasts. Segmented volumes of the Δ*chlL* dark cell (**a**) and Δ*chlL* 1h cell (**b**). (i) Raw slice; (ii) Raw slice superimposed with the segmentation; (iii) Segmented volume of the cell. Cellular components are labelled and coloured accordingly. Nascent thylakoid: NT; chloroplast envelope: CE; plasma membrane: PM; nuclear envelope: NE; starch granule: SG; and mitochondria: Mito; plastoglobule: PG. Scale bars = 1 µm. **c**, Normalized percentage of segmented chloroplast membrane contents of Δ*chlL* in dark and Δ*chlL* 1h was labelled on segmented volume. CE serves as the reference. For the dark conditions, 3,465,807 (1,382,638 for NT; 2,083,169 for CE) voxels are included. For the 1h condition, 11,453,730 voxels (7,827,824 for NT; 3,625,906 for CE) are included.

To investigate the distribution of protein complexes during thylakoid biogenesis, we conducted super-resolution live-cell fluorescence microscopy imaging of Δ*chlL* cells during the greening process. The Chl fluorescence signal observed in cells predominantly arises from active PSII centers and thus provides a direct readout of PSII synthesis and distribution in thylakoid membranes ^31^. A weak Chl signal was detected in the cells at the early stage of greening (0-2 h; Fig. 3a), consistent with the measured reduction in Chl levels (Fig. 1c). Intriguingly, super-resolution imaging and fluorescence profile analysis revealed that Chl fluorescence became detectable at 1 h, distributed throughout the cup-shaped chloroplast envelope, instead of being limited to the T-zone near the pyrenoid (as indicated by EPYC1-Venus using the pLM017 plasmid ^32,33^) (Fig. 3a). These results suggest that PSII biosynthesis and assembly occurred at multiple locations across the entire chloroplast, rather than been restricted to a single T-zone ^14,20^.

**Fig. 3.**
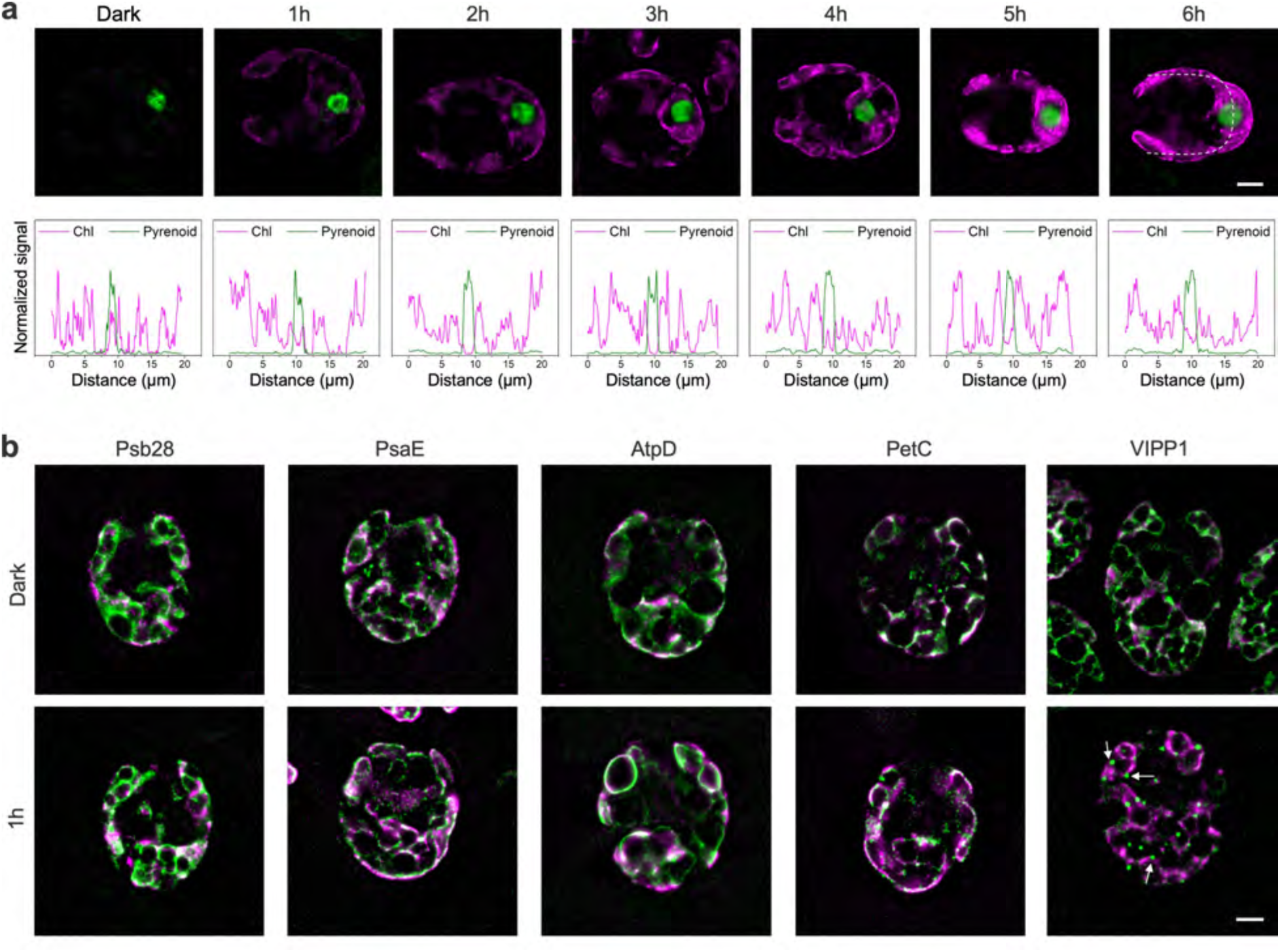
Live-cell super-resolution imaging reveals decentralized and globalized thylakoid biogenesis. **a**, Super-resolution fluorescence images of Δ*chlL* EPYC1-Venus cells during the greening process (top). Chl fluorescence is shown in magenta and Venus fluorescence in green (applies to all subsequent panels). Signal intensity profiles (bottom) were obtained by measuring the fluorescence intensity along the U-shaped arrow across the pyrenoid (shown in the image at 6 h), illustrating Chl (magenta) distribution during thylakoid biogenesis. Images at different time points represent different cells with similar chloroplast structures. Thirteen phase images were acquired per cell for SIM^2^ reconstruction. Scale bar = 2 µm. **b**, Representative merged fluorescence images of Δ*chlL* Psb28-Venus, Δ*chlL* PsaE-Venus, Δ*chlL* AtpD-Venus, Δ*chlL* PetC-Venus, and Δ*chlL* VIPP1-Venus strains. VIPP1 puncta are marked with white arrows. The brightness of both Chl autofluorescence and Venus fluorescence signals was adjusted to better reveal relative localization rather than absolute abundance. Scale bar = 2 µm.

Vesicle-inducing protein in plastids 1 (VIPP1, also named IM30) is vital for the biogenesis and maintenance of thylakoid membranes ^34–36^, assembling in higher-ordered structures on flat and gently curved membranes to deliver lipids for thylakoid membrane development and remodelling ^37^. At 0 h, VIPP1-Venus was broadly distributed in the chloroplast stroma (Fig. 3b, Fig. S5). After 1 h of illumination, it formed puncta dispersed across the chloroplast (Fig. S5), resembling the light-induced *in vivo* VIPP1 puncta observed in cyanobacteria ^38^. These changes are consistent with a dynamic redistribution of VIPP1 during early greening, although the fluorescence data alone do not establish the molecular mechanism of VIPP1 action. At 2-6 h, VIPP1 puncta became less prominent, and VIPP1 increasingly co-localized with the newly formed thylakoid membranes (Fig. S5). Overall, the changes in VIPP1 suggested that the biogenic sites of thylakoid membranes were distributed throughout the chloroplast in coordination with the redistribution of VIPP1, and that this spatial organization is tightly regulated in response to developmental or environmental cues, indicating the multiple roles of VIPP1 in membrane formation, maintenance, and remodelling ^39^.

To further examine the localization of various photosynthetic complexes, we generated constructs expressing Psb28-Venus, CP26-Venus, PsaE-Venus, PerC-Venus, AtpC-Venus, AtpD-Venus, Lhca1-Venus, and Lhca3-Venus in Δ*chlL*. Open reading frames (ORFs) were amplified by PCR from genomic DNA and cloned in frame with a C-terminal Venus YFP, driven by the strong PsaD promoter ^32,33,40,41^. The Venus-tagged proteins were correctly transported into the chloroplast (Fig. 3) and fluorescence tagging did not result in any detectable effects on cell growth or greening. Psb28 transiently interacts with PSII assembly complexes and PSII monomers, playing a role as an assembly factor in the assembly and repair of PSII ^42–44^, and CP26 (LHCB5) is a monomeric antenna protein of PSII. At 1 h, both proteins exhibited a global distribution throughout the chloroplast, consistent with the intrinsic Chl signal assigned to PSII (Figs. 3a, 3b, Fig. S6). PsaE is a peripheral subunit that binds to the PSI core ^45,46^, and Lhca1/Lhca3 serve as the light-harvesting antenna subunits associated with the PSI core ^45^. Similar to the PSII subunits, these proteins also exhibited a chloroplast-wide distribution, indicating the global biosynthesis of PSI complexes (Fig. 3b, Fig. S6). Likewise, fluorescence imaging of AtpC-Venus, AtpD-Venus, and PetC-Venus cells at 1 h revealed global biosynthesis of both F-ATPase and Cyt *b*_6_*f* complexes in the chloroplast (Fig. 3b, Fig. S6). Collectively, these results support the conclusion that the biosynthesis of photosynthetic complexes embedded in thylakoid membranes take place throughout the whole chloroplast along with the membrane recovery, highlighting a highly coordinated global biogenesis of thylakoid membranes.

### In-cell cryo-ET unveils morphological transformation of thylakoid membranes

To dissect the native molecular architecture of thylakoid membranes at the early stage of the greening process, we employed cryo-ET coupled with cryo-FIB on Δ*chlL* cells. The reconstructed tomograms, combined with segmented 3D volumes, provided native views on the change in thylakoid membrane content and morphology within Δ*chlL* cells during greening, revealing unprecedented molecular details (Fig. 4, Figs. S7, S8, Table S2). In agreement with our thin-section TEM and cryo-FIB/SEM volume data, in Δ*chlL* at 0 h, NTs (or residual membranes) were sparsely distributed throughout the chloroplast and displayed a variable and fragmented morphology in the basal region, lobes, and the T-zone, whereas stacked thylakoid membranes were diminished (Fig. 4a-c top, Fig. S8). Most membrane protein complexes, except for F-ATPases, were barely detectable, and the thylakoid lumen became highly disordered and filled with small densities (Fig. S7). Notably, pyrenoid tubules remained structurally intact, whereas mini-tubules were diminished (Fig. 4d), consistent with a compositional distinction between pyrenoid tubules and canonical thylakoids, despite their continuity in the WT ^30^. In Δ*chlL* at 1 h, NTs were elongated and became more continuous in the basal region, lobes, and the T-zone, consistent with light-triggered membrane synthesis and expansion (Figs. 4a-c, bottom), and the initial stacking of multiple NTs could be readily observed (Fig. 4b, bottom). In parallel, the overall morphology of tubules within the pyrenoid of Δ*chlL* at 1 h remained similar to that of the Δ*chlL* at 0 h, but mini-tubules reemerged within the pyrenoid tubules (Fig. 4d). By 24 h, the thylakoid morphology was fully restored and indistinguishable from the WT (Fig. S8). In addition to the direct visualization, we also quantified the membrane content during the greening process, which showed a nearly 4-fold change during the first hour of light treatment and nearly full restoration at 24 h (Fig. 4e).

**Fig. 4.**
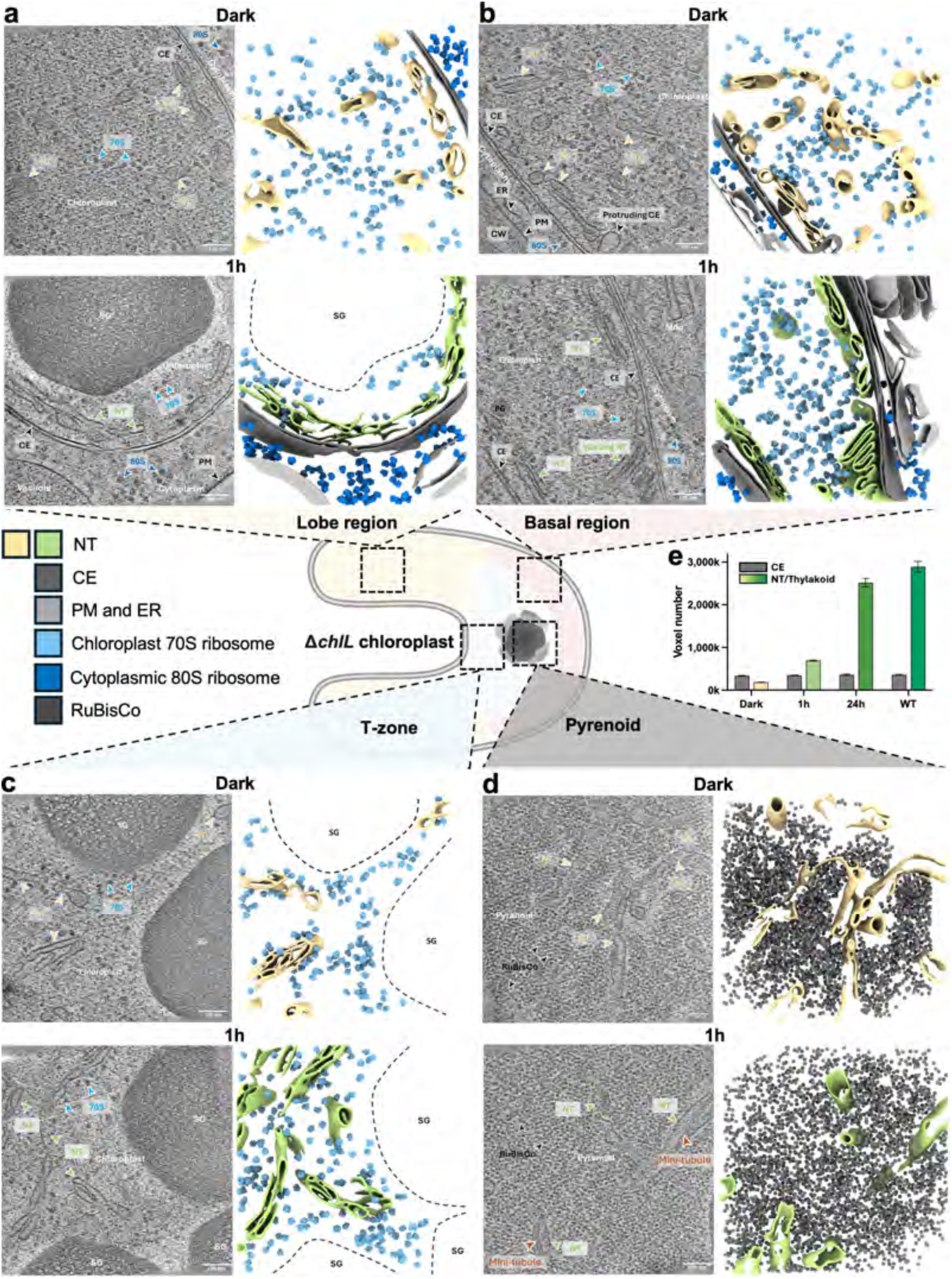
In-cell cryo-ET of incipient thylakoid biogenesis across the chloroplast. **a-c**, Representative tomographic slices (left) and segmented volumes (right) of Nascent thylakoids in the lobe (**a**), basal region (**b**), and T-zone (**c**) of Δ*chlL* chloroplast in dark (top) and after 1 h treatment (bottom). SG region is indicated by the dashed line in segmented volumes. **d**, A representative tomographic slice and segmented volume of pyrenoid of Δ*chlL* chloroplast. A schematic of the chloroplast and corresponding regions are show in the middle. Cellular components are coloured and labelled accordingly, as indicated in the schematic. Nascent thylakoid: NT, chloroplast envelope: CE, plasma membrane: PM, cell wall: CW, chloroplast 70S ribosomes: 70S, cytoplasmic 80S ribosomes: 80S, endoplasmic reticulum: ER, starch granule: SG, and mitochondria: Mito. **e**, A bar chart showing the voxel number of segmented membrane content across four conditions. *n* of dark-treated Δ*chlL* = 9, *n* of 1-hour-light-treated Δ*chlL* = 7, *n* of 24-hour-light-treated Δ*chlL* and WT = 3. One-way ANOVA is applied, *P* < 0.0001.

To further characterize the morphological transformation from NTs to mature thylakoids, the curvature of the NTs in Δ*chlL* cells was measured and compared with that of standard mature thylakoids in the WT. Remarkably, NTs appeared more curved and variable than those in the mature thylakoids (Figs. 5a, 5b). During greening, the curvature of the NTs remarkably decreased with light treatment, coupled with a continuous decrease in the variance (Figs. 5a, 5b). In parallel, the lumen space of the NTs also decreased during light treatment. (Figs. 5a, 5c). The mean intermembrane distance of mature thylakoids in WT cells and Δ*chlL* at 24 h measured 8.00 ± 1.60 nm and 7.6 ± 1.56 nm, respectively, similar to the value reported in previous studies ^47^, whereas the mean intermembrane distance of NTs in the Δ*chlL* cells at 0 h and 1 h measured 24.97 ± 12.01 nm and 17.32 ± 4.45 nm, respectively. Of note, the lumen density also decreased dramatically after light treatment (Fig. 5d), likely reflecting the coupling of lumenal protein dynamics with thylakoid biogenesis. Taken together, our cryo-ET data show a pronounced morphological transformation of NTs during the greening process and highlight the distinctive features of NTs at the molecular level in a spatiotemporal manner.

**Fig. 5.**
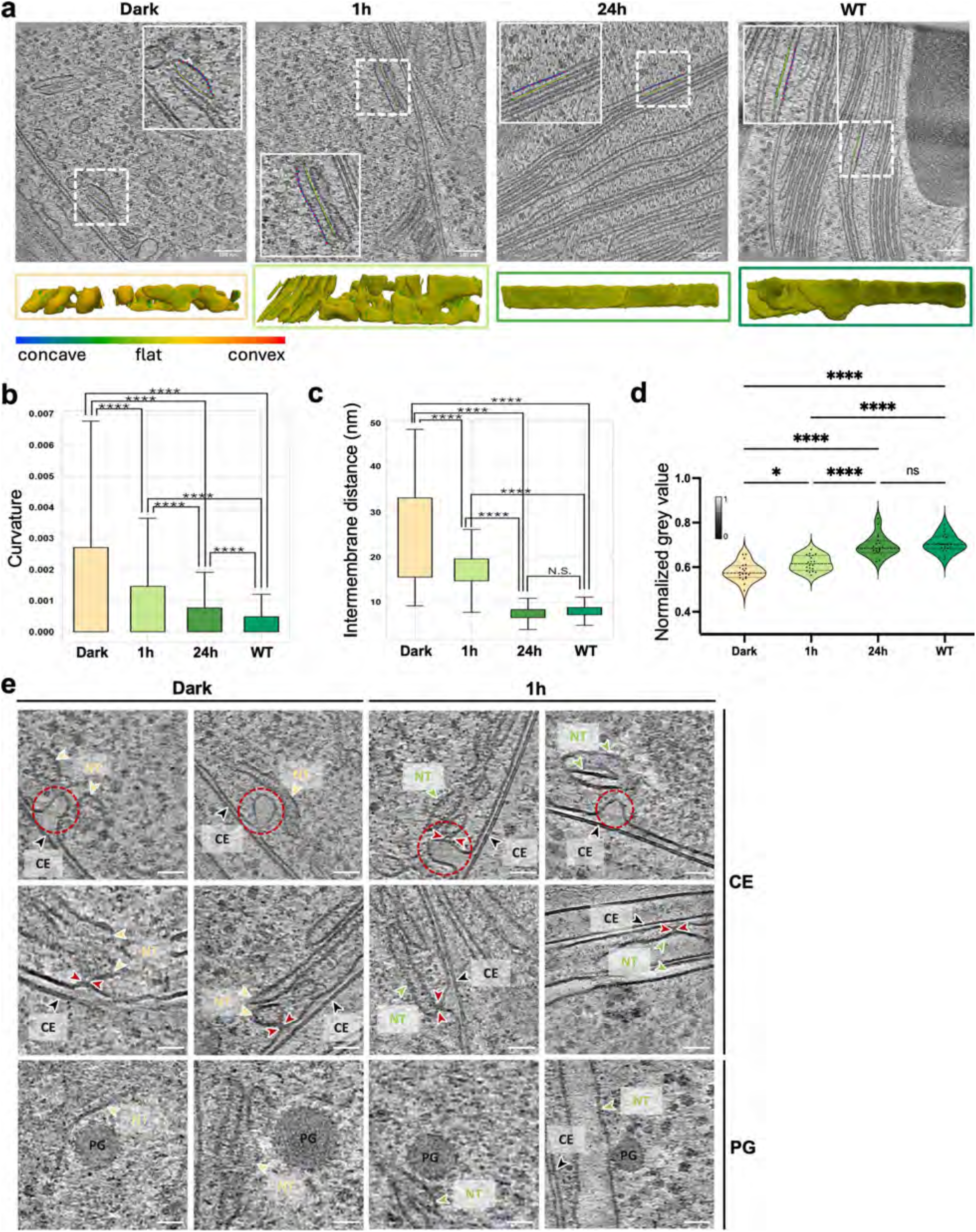
In-cell analyses of membrane remodelling in thylakoid biogenesis. **a**, Representative tomographic slices (top panel) with segmented volumes (bottom panel) exemplifying the measurement of lumenal intermembrane distance and density of thylakoids and surface curvature of thylakoids in different conditions. The distance and density measurements are taken between the blue and yellow lines indicated in inset. Curvature measurement is based on the segmented volumes generated using the same parameter, membrane fridges on the boundary of tomograms are excluded from the measurement. **b**, Surface curvature of thylakoids in different conditions. Dark-treated Δ*chlL*: 1.70 × 10^−3^ ± 1.82 × 10^−3^, SE = 1.41 × 10^−6^, 1 h Δ*chlL*: 1.02 × 10^−3^ ± 1.32 × 10^−3^, SE = 0.597 × 10^−6^, 24 h Δ*chlL*: 0.55 × 10^−3^ ± 0.71 × 10^−3^, SE = 0.258 × 10^−6^, and WT: 0.31 × 10^−3^ ± 0.36 × 10^−3^, SE = 0.122 × 10^−6^. One-way ANOVA test is applied, **** = *P* < 0.0001, *n* of tomograms for Δ*chlL* in dark, with 1 h treatment, 24 h treatment, and WT are 9, 7, 3, and 3, respectively. **c**, Lumenal intermembrane distances in different conditions. The measurement is exemplified in (**a**) by assessing the distance between the paired spots on the blue and yellow lines along the membrane. Dark-treated Δ*chlL*: 24.97 ± 12.01 nm, SE = 0.671, 1 h Δ*chlL*: 17.32 ± 4.45 nm, SE = 0.249, 24 h Δ*chlL*: 7.47 ± 1.56 nm, SE = 0.087, and WT: 7.99 ± 1.60 nm, SE = 0.089. One-way ANOVA test is applied, **** = *P* < 0.0001, ns = no significance, *n* of each condition = 320. **d**, Grey value measured for the lumenal space of thylakoids in different conditions. Dark-treated Δ*chlL*: 0.581 ± 0.041, SE = 0.009, 1 h Δ*chlL*: 0.618 ± 0.034, SE = 0.008, 24 h Δ*chlL*: 0.696 ± 0.048, SE = 0.011, and WT: 0.707 ± 0.040, SE = 0.009. One-way ANOVA test is applied, * = *p* = 0.0269 < 0.05, **** = *P* < 0.0001, ns = no significance, *n* = 20 for each sample. **e**, Representative tomographic slices showing interactions between NTs and chloroplast envelop and plastoglobules in Δ*chlL* cells in dark and 1 h. NT and CE are labelled accordingly, the protruding CE is indicated in red dotted circles, and contacting points are indicated by red arrowheads. Nascent thylakoid: NT, chloroplast envelope: CE, and plastoglobule: PG. Scale bar = 50 nm.

### Invagination of chloroplast inner envelop membrane potentially supplies lipids for thylakoid biogenesis

In the tomograms, we frequently observed that the IEM of chloroplasts periodically extended inward toward the NTs (Fig. 5e, top two rows). Similar IEM invaginations have also been observed in WT *C. reinhardtii* with mature thylakoid membranes ^30^. Notably, the lumen density of the invaginated IEMs was much lower than that of the NTs detected at 0 h, suggesting that NTs were not directly formed through the budding of IEMs (Fig. 5e, top two rows). Instead, we observed direct contact between the IEM and NTs (Fig. 5e, red arrowheads), which may be crucial for the transfer of lipids and proteins during thylakoid biogenesis. Additionally, NTs situated away from the IEM were frequently associated with plastoglobules (PGs), which are responsible for maintaining carotenoid and lipid homeostasis in thylakoids ^30,48^ (Figs. 3c, 3d, 5e). The PGs varied in size and displayed distinct surface densities (Fig. 5e, bottom). In dark-treated Δ*chlL* cells, PGs were found in 11 cases, all closely associated with NTs. In Δ*chlL* cells at 1 h after illumination, PGs were identified in 24 cases, with 22 closely associated with NTs. These findings suggest the possibility that PGs act as reservoirs of lipids and biosynthetic enzymes during thylakoid biogenesis ^48,49^.

### Global translation by chloroplast 70S ribosomes orchestrates early thylakoid biogenesis

Given that thylakoid biogenesis entails extensive protein synthesis, our fluorescence imaging data also suggest that this process coincides with the assembly of a variety of protein complexes. We then assessed *in situ* translation activity in the chloroplast by examining the distribution of chloroplast 70S ribosomes during the biogenesis of thylakoids. We first conducted template matching on the 70S ribosomes in the chloroplast across the three conditions: Δ*chlL* in the dark (dark), Δ*chlL* receiving 1-hour light treatment (1h), and the WT in the dark (Figs. S9, S10c). Through extensive classification and structural refinement of the picked 70S ribosome particles, we obtained sufficient resolution to identify two major classes of 70S ribosomes in Δ*chlL* cells (Fig. S9d, Table S3): one exhibiting a prominent mRNA density at the P-site in the center of the 70S ribosome, corresponding to actively translating ribosomes ^50,51^, and another lacking this density, corresponding to non-translating ribosomes (Fig. 6a, Figs. S9a, S9b). In contrast, only a single class with a clear mRNA density at the P-site was resolved in WT cells under dark conditions (Fig. 6a, Fig. S9c). Notably, only 20.4% of 70S ribosomes in Δ*chlL* cells were engaged in translation at 0 h, whereas the proportion of actively translating ribosomes increased to 83.0% after 1 h light exposure (Fig. 6a). In comparison, nearly all 70S ribosomes in WT chloroplasts remained in an active translation state regardless of light availability (Fig. 6a). We then questioned whether the translating 70S ribosomes were evenly distributed across the chloroplast or clustered in certain regions. Through the mapping back of each individual 70S ribosomes in the Δ*chlL* cells in the dark and light (Fig. 6b), we found that the translating 70S ribosomes were more enriched in the T-zone in the dark and became evenly distributed after 1 h light exposure (Figs. 6c, 6d). The higher level of translating 70S ribosomes near the T-zone in the dark may reflect localized protein synthesis associated with the pyrenoid (e.g. RuBisCO, EPYC1, carbonic anhydrase) ^9^ or other T-Zone functions, but the precise role remains unclear; whereas, the even distribution after light treatment indicates the onset of global thylakoid biogenesis at the translational level.

**Fig. 6.**
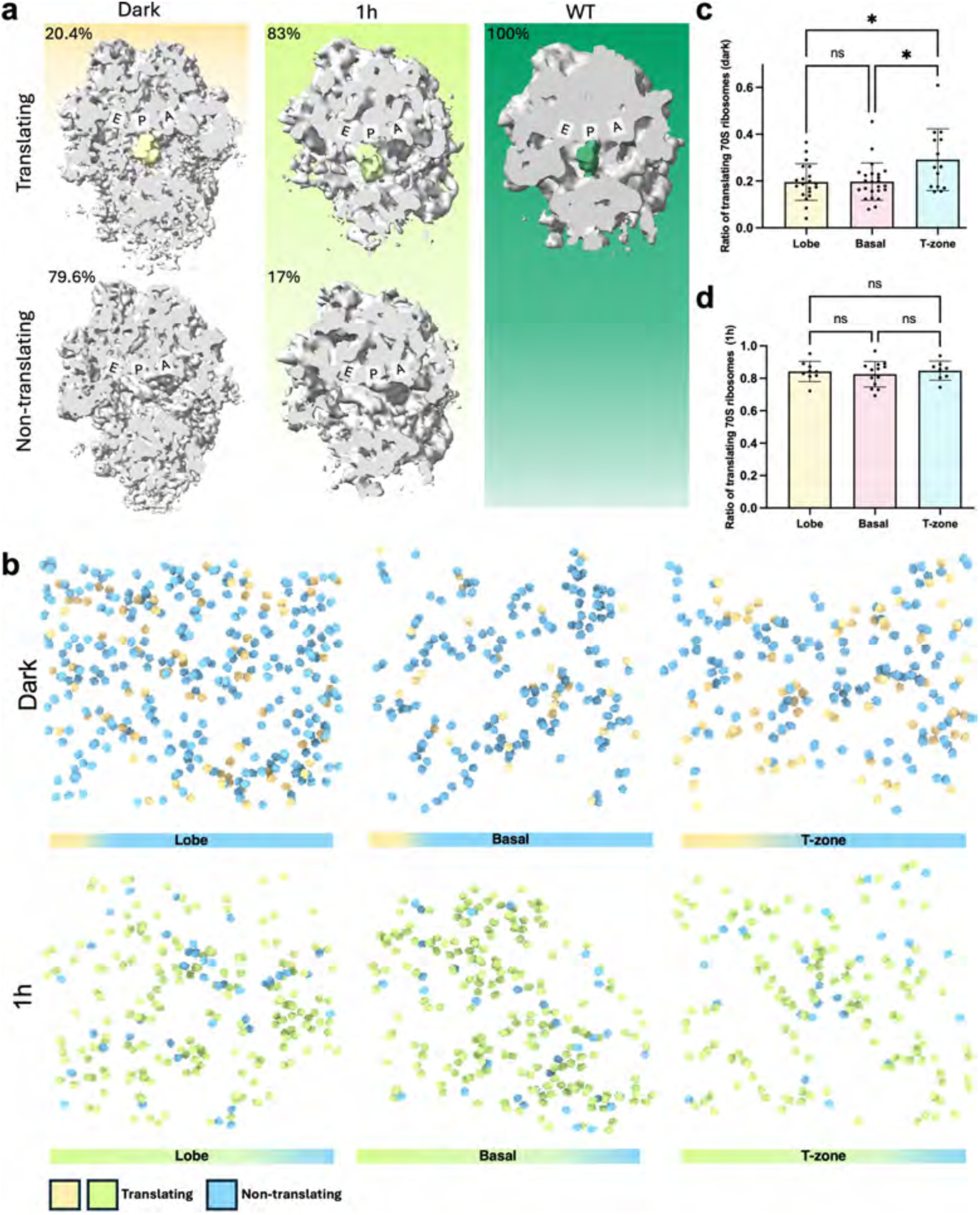
Translational activity of chloroplast 70S ribosomes at the early stage of thylakoid biogenesis. **a**, In-cell structure of chloroplast 70S ribosome in translation/non-translation states in three conditions: Δ*chlL* dark, Δ*chlL* 1 h, and WT in dark. For better visualization, the t-RNAs occupying the P site is low-pass filtered and coloured accordingly, and the main body of ribosome is coloured grey. **b**, Mapping back of translating and non-translating chloroplast 70S ribosomes in different regions of chloroplasts in Δ*chlL* dark and 1 h. Non-translating ribosomes are coloured light blue, translating ribosomes are coloured light yellow in dark and light green in 1 h, respectively. The gradient bar indicates the abundance of ribosome population. **c-d**, Bar charts showing the proportion of translating and non-translating chloroplast 70S ribosomes in different regions of chloroplasts of Δ*chlL* dark (**c**) and Δ*chlL* 1 h (**d**). One-way ANOVA test is applied, for Δ*chlL* in dark, *n* of lobe = 20, mean = 0.196 ± 0.078, SE = 0.018; *n* of basal = 24, mean = 0.198 ± 0.080, SE = 0.016; *n* of T-zone = 14, mean =0.291 ± 0.132, SE = 0.035. For Δ*chlL* 1 h, *n* of lobe = 9, mean = 0.842 ± 0.063, SE = 0.021; *n* of basal = 13, mean = 0.825 ± 0.079, SE = 0.022; *n* of T-zone = 8, mean = 0.847 ± 0.059, SE = 0.021. * = *p* < 0.05, ns = no significance.

### Hierarchical synthesis and assembly of photosynthetic complexes during thylakoid biogenesis

Thylakoid membrane biogenesis and formation of the photosynthetic apparatus require the synthesis and assembly of major photosynthetic complexes such as PSI, PSII, Cyt *b*_6_*f*, and F-ATPase. To investigate the *de novo* biosynthesis of these photosynthetic complexes during the greening process, we used label-free mass spectrometry to quantify the protein abundance in Δ*chlL* cells collected at successive stages of thylakoid biogenesis. The abundance of each protein was normalized to its maximum abundance under equal protein loading to obtain its accumulation trajectory (Figs. 7a, 7b, Fig. S11, Table S4).

**Fig. 7.**
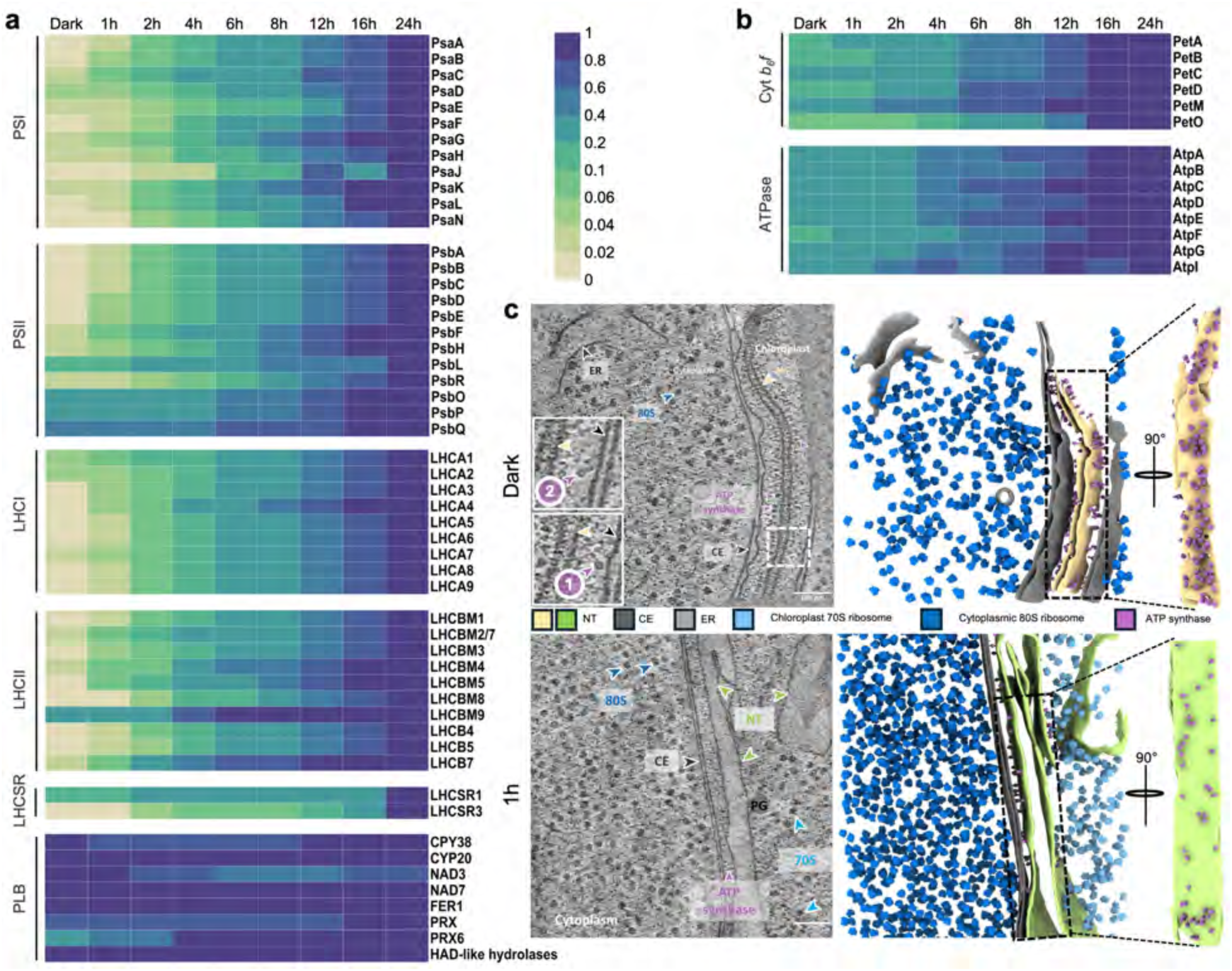
Protein synthesis and recovery during thylakoid biogenesis. **a-b**, Quantification of normalized protein content based on mass spectrometry. PLB, prolamellar body. Whole-cell protein extracts were collected from Δ*chlL* cells undergoing thylakoid biogenesis. Data represent relative protein abundance; the relative abundance of each protein was normalized to its maximum abundance across at least three biologically independent replicates. **c**, Representative tomographic slice (left) and segmented volume (right) featuring the F-ATPase storage membrane (top) and NT after 1-hour light treatment (bottom), two zoomed-in views of F-ATPases are depicted in the insets of the top tomographic slice. NT, CE, 70S, 80S, F-ATPase, and ER are labelled accordingly. Membrane face-on views were shown on the side to illustrate the distribution of F-ATPases.

PSII and PSI subunits were present at relatively low levels at the early stages of thylakoid biogenesis (Fig. 7a). For Δ*chlL* at 0 h, PSI subunits were present at only 1–5% of their abundance levels at 24 h. After 1 h of the light treatment, the contents of PsaA, PsaB, and PsaC increased by 244%, 355% and 99%, respectively, indicating active *de novo* assembly of the PSI reaction center (RC) (Fig. S11a). This is consistent with the retention of 50%-70% of PSI assembly factors (e.g. Ycf3, Ycf4, Y3IP1) ^46^ in dark-adapted cells (Fig. S11a), and suggests that the early stages of PSI assembly are poised to proceed upon illumination. The PSII core components began to accumulate from the basal levels (∼1–2%) at 0 h (Fig. 7a). PsbA (D1) and PsbDEF (D2-cyt b559), which are essential for initial PSII RC assembly, increased by ∼200% after 1 h light treatment (Fig. S11a). PsbB and PsbC, required for the assembly of RC47 and the PSII core, exhibited rapid accumulation rates of ∼120–140% per hour during the first 2 h of greening. PsbR and PsbH, linked to RC47b synthesis ^1,43,52^, maintained high accumulation rates until 12 h and then reached stable levels (Fig. S11a). Notably, the lumen-localized oxygen-evolving complex (OEC) subunits of PSII, including PsbO, PsbP, and PsbQ, were detected with an initial level of 30–50% at 0 h relative to the 24 h level (Fig. 7a), which may contribute to the greater lumen density of NTs (Fig. 5e). These subunits, which lack a pigment-binding capacity, may not be tightly regulated by Chl availability ^53–55^.

In *C. reinhardtii*, the PSI–LHCI supercomplex contains up to ten peripheral antenna LHCI subunits associated with the PSI core ^56,57^, and LHCIIs include nine members, designated LHCBM1-9 ^58^. Additionally, light-harvesting complex stress-related proteins (LHCSRs) are active in the nonphotochemical quenching (NPQ) of photosystems ^59–61^. Our results showed that the major LHC subunits were present at low levels at the early stages of thylakoid membranes and were synthesized rapidly in response to illumination, in agreement with the biosynthesis profiles of PSI and PSII (Fig. 7a). Notably, LHCBM9 displayed relatively higher levels early in light-induced thylakoid biogenesis, highlighting the potential photoprotective role of LHCBM9 in facilitating light energy dissipation and preventing cell damage under stress conditions ^62^.

Unlike the low levels of PSI, PSII, and LHC proteins, F-ATPases (as indicated by the contents of AtpA-G and AtpI) were present in the dark (0 h) at approximately 20% relative to the 24 h level (Fig. 7b). The abundance of F-ATPases increased progressively throughout the greening process (from 1 h to 24 h). A similar pattern was observed for Cyt *b*₆*f* complexes, with ∼20% of PetA-D&M subunits preserved at 0 h relative to 24 h (Fig. 7b). In contrast, PetO, previously designated as subunit V of Cyt *b*₆*f* but appears to be involved in the cyclic electron cycle ^63,64^, displayed a slightly different synthesis profile that was not fully synchronized with the other of Cyt *b*₆*f* subunits.

We then investigated whether these complexes could be directly visualized in the tomograms of Δ*chlL* cells during greening. Except for F-ATPase, the PSI, PSII, and Cyt *b*_6_*f* complexes were not clearly detected either by direct observation or through template matching at 0 h and 1 h, but became evident at 24 h and in WT cells (Fig. S7a). Intriguingly, in Δ*chlL* at 0 h, some NTs were densely packed with F-ATPases (Fig. 7c), as verified by template matching and subtomogram averaging (Figs. S10a, S10b), suggesting that these membranes act as reservoirs for F-ATPases. We therefore refer to these special membranes as “F-ATPase storage membranes”. Strikingly, the F-ATPase density in the storage membranes reached ∼2,200 particles/μm^2^, far exceeding the ∼700 particles/μm^2^ observed with the NTs at 1 h (Fig. 7c, also see the Methods section) and higher than the ∼1,700 particles/μm² reported for thylakoids ^47^. In addition, some F-ATPase structures were observed floating in the stroma (Fig. S7b), which might represent assembly intermediates to be incorporated into thylakoids ^65^.

These F-ATPase storage membranes are reminiscent of the prolamellar bodies (PLB) observed in plants, which contain ATPases, Cyt *b*_6_*f*, and OEC proteins and serve as precursors to chloroplast thylakoids during light-induced greening ^66–69^. Our proteomic data also revealed that some proteins reported in the PLB ^67^, such as protochlorophyllide reductase (POR), thioredoxin-dependent peroxiredoxin (PRX), NADH dehydrogenase subunit 3 (NAD3), and high chlorophyll fluorescence (HCF136), were highly abundant in the 0 h samples (Fig. 7a, Fig. S11b).

Most factors involved in photosystem biosynthesis and repair were detected at the abundance levels of 30%-90% at the early stages of thylakoid biogenesis compared to the levels at 24 h (Fig. S11b), substantially higher than the aforementioned PSI, PSII, and LHC proteins. This suggests that they are required for rapid initiation of photosynthetic apparatus assembly and thylakoid membrane development. Among them, the LHC-like protein MSF1 was reported to be a factor for PSI maintenance, showing an accumulation pattern resembling that of Chl biosynthesis ^70^. The PSII biogenesis factor PsbN, also known as photosystem biogenesis factor 1 (PBF1) ^71,72^, displayed a high abundance at 0 h compared to that at 24 h. However, its content declined rapidly at the early stage and subsequently recovered, which may imply a shift in its function from promoting RC formation to PSII photoprotection ^1,43^. PsbP-binding protein 1 (PPD1), a PSI assembly protein involved in the initial assembly steps of the PSI RC from the lumenal side ^16,46,73^, also exhibited a relatively high level at the beginning of greening, followed by the reduction in abundance as thylakoid development progressed. At 0 h, CURVATURE THYLAKOID1 (CURT1) accumulated to approximately 50% of its abundance at 24 h, which may account for the pronounced membrane curvature of NTs at the earlier stages of biogenesis ^74,75^ (Fig. 5b, Fig. S11b). The constant presence of chloroplast envelope-localized lipid synthases MGDG1 and DGDG1 ^5^ and fibrillins for lipids transport ^76^ throughout the greening process suggests active lipid synthesis during thylakoid membrane development (Fig. S11b).

Moreover, proteins associated with the electron transport chain (ferredoxin, plastocyanin, plastid terminal oxidase (PTOX)), carbon metabolism (Calvin–Benson cycle, respiration, photorespiration), gene expression (transcription, translation), and the TOC–TIC translocon system responsible for protein transport from the cytoplasm ^77^ were relatively abundant at the early stages of greening (Fig. S11b). In contrast, some CO_2_-concentrating mechanism proteins, such as CAH1, LCIA, HLA3, and BST1, accumulated more slowly before 24 h.

Collectively, the proteomic results revealed that photosynthetic protein components accumulate to various levels at different stages of thylakoid biogenesis, indicating a hierarchical assembly of photosynthetic complexes and the distinct roles of individual protein components in supercomplex assembly and the regulation of photosynthetic performance (e.g. light transduction, photoprotection) during thylakoid development. In addition, the storage membranes, potentially representing a primitive form of PLB, may serve as repositories of protein complexes and other factors essential for subsequent thylakoid biogenesis.

### Stepwise assembly of functional photosynthetic supercomplexes during thylakoid biogenesis

To explore the assembly of functional photosynthetic supercomplexes during thylakoid biogenesis, we determined the assembly status of photosynthetic complexes in isolated thylakoid membranes using blue native-polyacrylamide gel electrophoresis (BN-PAGE) combined with immunoblot analysis (Fig. 8a).

**Fig. 8.**
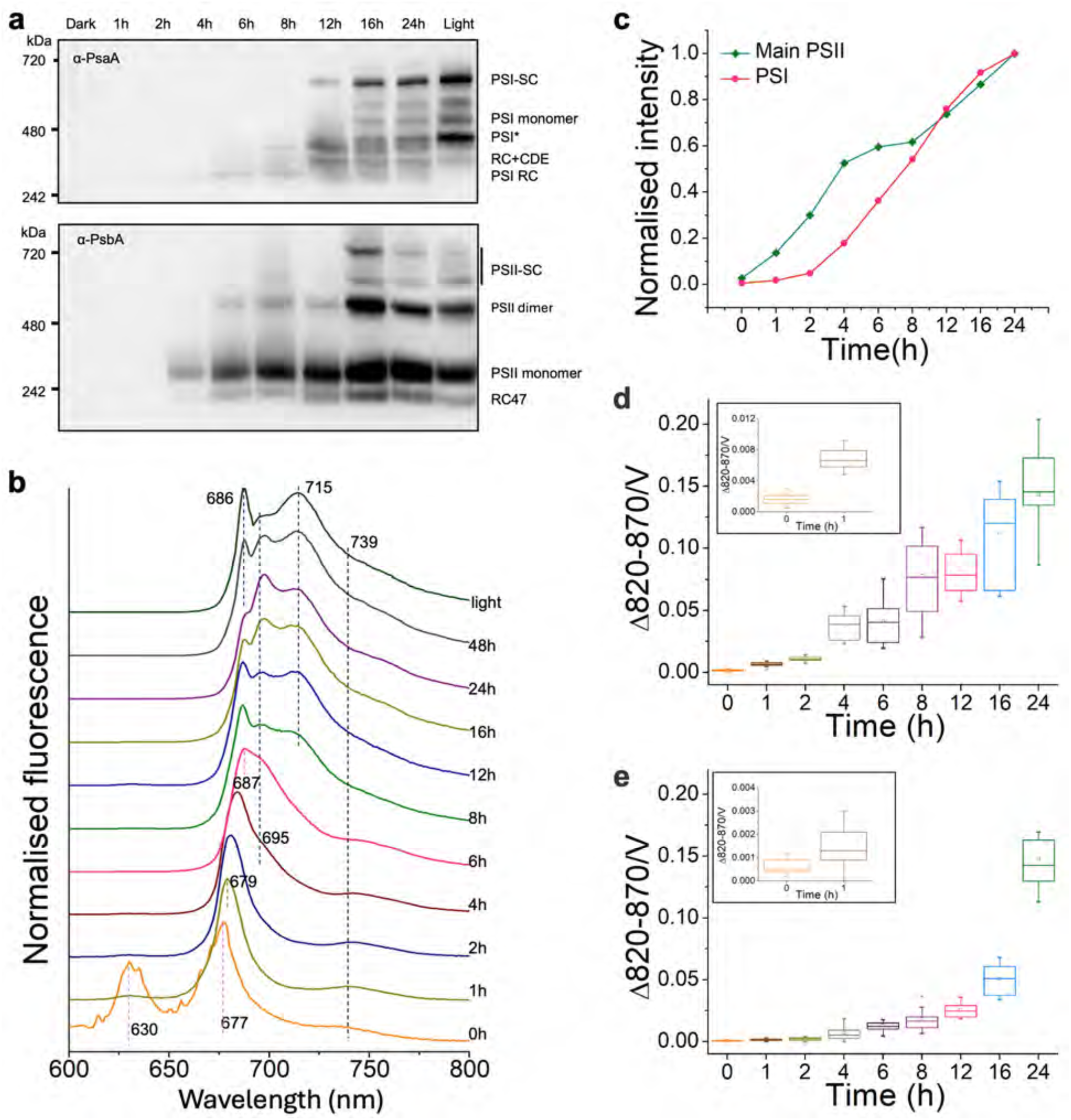
Stepwise assembly of photosynthetic supercomplexes. **a**, Blue-native-PAGE and immunoblot analysis of thylakoid membranes during thylakoid biogenesis using α-PsaA and α-PsbA antibodies. The intermediate PSI complex is annotated based on ^45^, and PSI* that lacks PsaG and PsaK is labelled according to ^106^. **b**, 77K fluorescence emission spectra measured from equal numbers of cells. All spectra were normalized to the maximum intensity. The emission peak at 630 nm corresponds to free chlorophyllide, peaks at 677-695 nm represent PSII, and peaks at 739 nm and 715 nm are attributed to PSI. **c**, 77K fluorescence peak intensities associated with PSII and PSI were quantified and normalized to their respective maximum value. **d**, P700 slow kinetics measured by PAM-KLAS system. The absorbance changes between 820 nm and 870 nm induced by a saturating pulse under dark conditions represent the maximal P700 pool size (*n* > 12). **e**, P700 slow kinetics measured by PAM-KLAS system under far-red light. Absorbance changes at 820-870 nm reflect the P700 pool involved in active cyclic electron transport (*n* > 12).

During thylakoid biogenesis, PSI RC became detectable at 4 h, followed by the formation of higher-ordered PSI assemblies, PSI monomers, and supercomplexes (PSI-SC). Both RC47 intermediates (or CP43-free PSII monomer) and PSII monomers were also detected at 4 h, prior to the detection of PSII dimers and PSII-SC. Interestingly, the RC47 content increased in parallel with the accumulation of PSII monomers, dimers, and PSII-SC, suggesting active PSII assembly and repair due to assembly lines increasing over time with growing membrane surfaces.

Furthermore, we recorded 77K fluorescence emission spectra (excited at 435 nm) of the Δ*chlL* cells collected at different timepoints (Figs. 8b, 8c). The fluorescence emission at 0 h displayed a peak at 630 nm, corresponding to free protochlorophyllide ^78^, and a peak at 677 nm, attributed to the probable presence of unassembled Chl-containing photosynthetic proteins. Fluorescence at 630 nm was greatly reduced after 1 h along with Chl biosynthesis, while the 677 nm fluorescence peak red-shifted to 679 nm (Cyt b559-D1D2 PSII RC) ^79^, 695 nm (PSII RC47), and 687 nm (PSII core with CP43) ^80–82^. Consistent with the BN-PAGE and immunoblot results (Fig. 8a), the fluorescence peaks of RC47 (695 nm) and PSII core with CP43 (687 nm) were detected at 4 h and 6 h, respectively. Additionally, the fluorescence peak at 739 nm representing assembled Lhca subunits ^81^ gradually increased, and the fluorescence band associated with mature PSI (715 nm) was detected at 8 h. Our analysis further revealed that PSII exhibited a faster accumulation than PSI at the early stages of thylakoid biogenesis (Fig. 8c), and Chl *b* was involved in the assembly of LHC proteins in PSII complexes at the early stage of thylakoid biogenesis (Fig. S12), consistent with its biosynthesis initiation at 1 h (Fig. 1c). Together, these results are consistent with a stepwise, hierarchical assembly of PSI-SC and PSII-SC during greening, suggesting a conserved mechanism governing photosynthetic complex formation across diverse photoautotrophs ^83,84^.

In addition, P700 oxidation kinetics were monitored to delineate the assembly of functional PSI during thylakoid biogenesis (Fig. S13). A weak P700 oxidation signal was detected (Δ820nm-870nm) after 1 h illumination (Fig. 8d), followed by a progressive increase upon illumination, indicating ongoing PSI biogenesis, which is in agreement with our proteomic analyses (Figs. 7a, 8a, Fig. S11b). At 4-h illumination, a substantial enhancement in the maximum pool of P700 was detected (Fig. 8d), coinciding with elevated PSI RC content and PSII monomer accumulation (Fig. 8a), accompanied by an enhanced oxygen-evolution rate (Fig. S2d). The saturation pulse (SP)-induced Δ820nm-870nm signal under far-red illumination, which was used as a proxy for cyclic electron transport (CET) around PSI, only increased exponentially after 24-h illumination (Fig. 8e), coinciding with the accumulation of PetO (Fig. 7b) which has been linked to CET ^63^. These results, combined with the low abundance of CET-associated PGRL1 and PGR5 ^85^ at early thylakoid biogenesis stages (Fig. S11b), suggest linear electron transport predominates initially. A ∼4-8 h delay was observed between the increase in the maximum pool of P700 and the rise in CET around PSI, coinciding with concurrent enhancement of both the 739 nm and 715 nm emission peaks in 77K fluorescence spectra (Fig. 8b). Since electron transport-associated proteins were already present during early thylakoid biogenesis (Fig. S11b), the accelerated PSI reduction kinetics were likely due to enhanced electron transfer from PSII, arising from the progressive assembly of functional PSII complexes and the coordinated organization of both photosystems in the developing thylakoid membranes. Indeed, the formation of PSI and PSII supercomplexes is tightly interconnected ^52^. Moreover, our data show that PTOX accumulation coincided with early PSII biogenesis (Fig. S11b), suggesting that PTOX temporarily provides an alternative electron sink downstream of PSII, functionally compensating for limited PSI availability and permitting PSII function before mature PSI is established at the early stages of thylakoid biogenesis ^86^.

## Discussion

The biogenesis of thylakoid membranes and assembly of photosynthetic complexes are fundamental processes underpinning oxygenic photosynthesis in plants, algae, and cyanobacteria ^5,6,8^. Despite substantial efforts in the past decades, the spatiotemporal dynamics and mechanisms governing these processes remain unclear. Some models proposed that photosystem biogenesis takes place at the T-zone close to the pyrenoid, where ribosomes, mRNAs, and nascent photosynthetic proteins accumulate ^14,15,20^, whereas earlier studies on the *y-1* mutant also showed membrane materials extended from IEM, suggesting envelope-associated thylakoid formation during greening ^19,21^. To date, there is no comprehensive theoretical framework that integrates the spatial and temporal dynamics of photosynthetic protein complex assembly with the formation of thylakoid lipid membranes, supported by molecular-level mechanistic evidence.

In this study, we established a light-triggered platform to modulate Chl synthesis and thylakoid development in the *C. reinhardtii* Δ*chlL* mutant. Upon growth in the dark, the Δ*chlL* strain exhibited a substantial reduction in thylakoid and Chl contents, whereas exposure to light for 24 h restored the structure and function of the thylakoid to those of the WT (Fig. 1, Figs. S8, S12b). This system enabled a systematic investigation of thylakoid membrane biogenesis and photosynthetic complex assembly during dark-to-light transitions with unprecedented resolution (Fig. 9). It is important to note that the greening process represents a physiologically relevant process rather than an experimental artifact, and a similar greening program naturally occurs during *Chlamydomonas* development. For example, during zygospore germination, dormant, pigment-deficient zygospores resume Chl biosynthesis and reconstruct functional thylakoid membranes as they differentiate into vegetative cells ^87^.

**Fig. 9.**
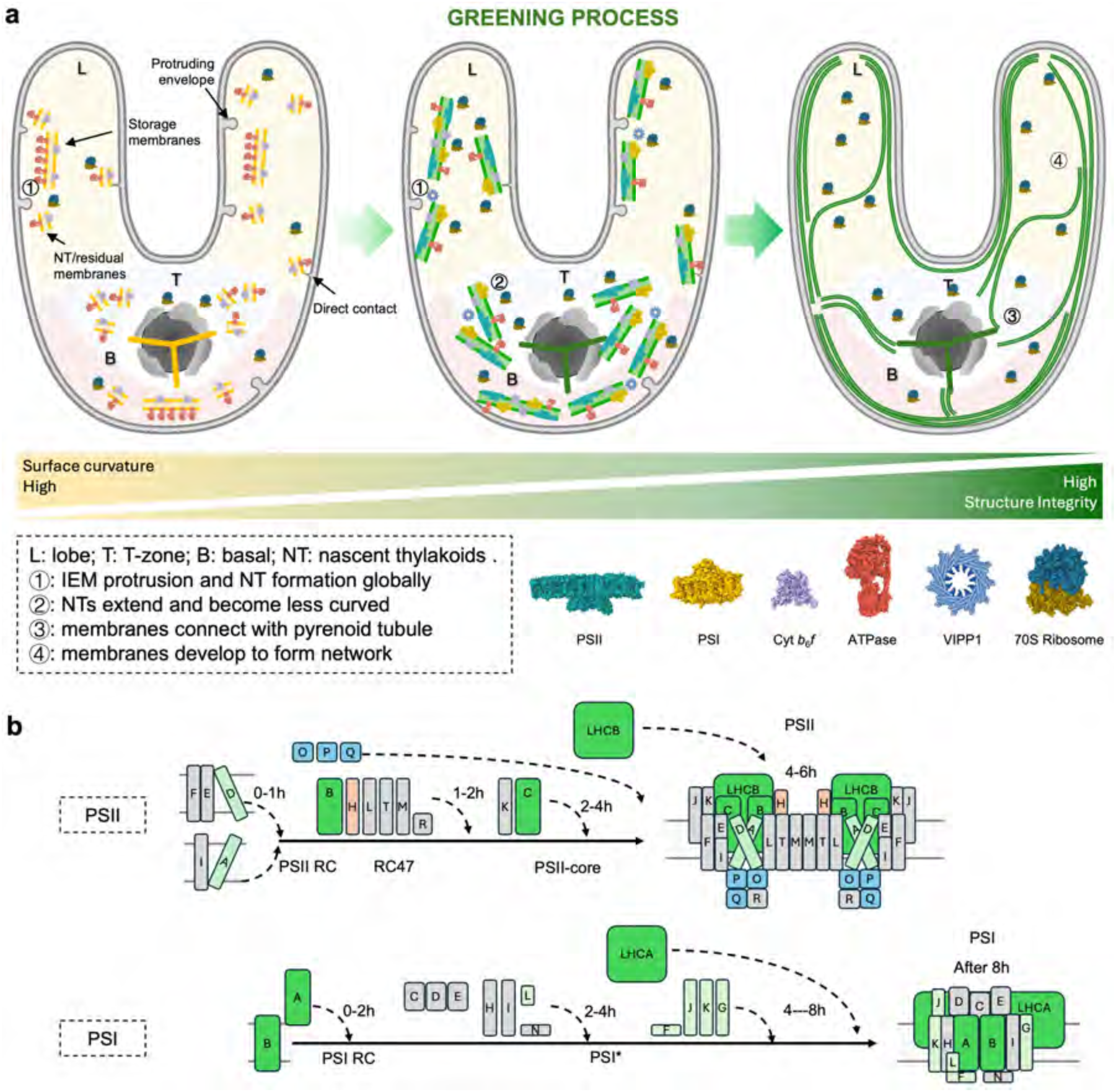
Schematic model of thylakoid biogenesis in *Chlamydomonas*. **a**, Chl-less membranes in the early dark phase are depicted in yellow; membranes undergoing biogenesis are shown in light green; fully reassembled membranes are in dark green. For clarity, protein complexes are not drawn in fully recovered chloroplast. The lobe region is coloured pale yellow, the T-zone light blue, and the basal region light pink. Note that the TM content, size, and relative distribution of complexes in the model are schematic and represent trends rather than absolute quantities. While full photosystems are displayed for illustration, stepwise assembly occurs progressively. VIPP1 puncta are represented by the oligomeric structure (PDB ID: 6ZVR) to reflect spatial localization inferred from fluorescence microscopy, but this does not imply that observed puncta are necessarily composed of oligomers. The ribosome structures shown in the model represent translating 70S ribosomes. Protein complex represented by structures from PDB as: PSII, 6KAF; PSI, 8H2U; F-ATPase, 6FKF; Cyt *b_6_f*, 2E74; VIPP1, 6ZVR; 70S Ribosome, 5MMM. Based on the electron microscopy data, events within the chloroplast are annotated according to location. Event ①: IEMs protrude towards the chloroplast stroma and NTs appear throughout the chloroplast, sometimes direct connections between IEM invaginations and NTs observed. Event ②: NTs expand with reduced curvature and intermembrane distance. Event ③: The developing and extended thylakoids establish the connection with pyrenoid tubules. Event ④: The developing thylakoid membrane fragments form a stacked, continuous network throughout the chloroplast. **b**, Stepwise assembly of photosystems during greening. The structure diagrams of PSII and PSI refer to 6KAF ^58^ and 6IJJ ^56^, respectively.

Using thin-section TEM, super-resolution fluorescence imaging, in-cell cryo-FIB/SEM volume imaging and cryo-ET, our time-resolved analyses show that during greening in the Δ*chlL* strain, thylakoid membrane formation and photosystem assembly occur at multiple chloroplast locations and are not confined to the T-zone region alone. Instead, at the early stage of thylakoid biogenesis, small fragmented thylakoid membranes (denoted NTs) and photosynthetic complexes displayed a globally distributed pattern throughout the entire chloroplast, including the basal region, lobes, and stroma adjacent to the pyrenoid (Fig. 9a). This interpretation is supported by the observed redistribution of translating 70S ribosomes throughout the chloroplast during early greening (Fig. 6). A globally distributed pattern may enhance the efficiency, robustness, and adaptability of thylakoid biogenesis, thereby enabling robust recovery of photosynthetic performance and environmental fitness. During 2-8 h illumination, the NTs gradually elongate and merge, exhibiting enhanced membrane stacking and increasing structural complexity. As the greening process progresses, the developing thylakoid membranes interconnect and fuse to form expanded membrane layers, which eventually mature into complete and functional thylakoid networks after 24-h illumination, with WT levels of protein and pigment abundance and photosynthetic activity (Fig. 9a).

Importantly, our observations extend classic *y-1* studies that also reported envelope-associated membrane emergence during greening. A large number of thylakoid biogenesis sites were detected near the chloroplast IEM, consistent with previous conclusions obtained with the *y-1* mutant ^19^. The IEM invagination in *C. reinhardtii*, as observed in this study (Fig. 5e) and previous work ^30^, may facilitate lipid transport from the IEM to thylakoid membranes to drive thylakoid formation and expansion ^5,88^. It is noteworthy that lipid transport does not occur through budding or fusion; as we observed, invaginated IEMs formed direct contact with NTs (Fig. 5e, red arrowheads). Additionally, during the initial stage of thylakoid biogenesis, PGs were often closely associated with thylakoid membranes (Figs. 4b, 5e), potentially serving as the sources of carotenoids and lipids for thylakoid formation. F-ATPase complexes were enriched in specific storage membranes (Fig. 7c), and PLB-containing proteins were present in high abundance (Fig. 7a). Given that chloroplast F-ATPase can modulate the proton motive force and regulate luminal pH under stress conditions ^89^, the presence of F-ATPase in the storage membranes raises the possibility that these membranes contribute to local proton gradient formation or maintenance. In this context, such proton motive force may support Tat-dependent protein translocation ^90^, which relies on ΔpH across the membrane. Direct mechanistic evidence is required to evaluate whether storage membranes are functionally competent in sustaining these processes.

IEM invagination and thylakoid-IEM contacts have also been commonly observed in land plants (such as *Arabidopsis*) with developing chloroplasts or under stress conditions or in cases where thylakoid development is inhibited ^83,91–93^. In C4 plants, the peripheral reticulum fills the space between the thylakoids and IEM, and a network bridge is constructed by budded vesicles ^94^. In cyanobacteria, thylakoid biogenesis sites have also been shown to involve multiple, spatially distributed membrane foci rather than a single nucleation site ^35,38,95,96^ and are close to the plasma membrane, although without direct fusion between the two adjacent membrane systems ^84,97^. The budding of the peripheral cytoplasmic membrane to drive the formation of chromatophore vesicles has been observed in purple photosynthetic bacteria (e.g. *Rhodobacter sphaeroides*) when shifting cell growth from microoxic to anoxic light conditions ^2,98,99^. Such membrane contacts may represent a universal mechanism for phototrophic organisms to facilitate the transfer of lipids, proteins, and other components between distinct membrane systems.

Recent studies have shown that VIPP1/IM30 can assemble into polymeric and higher-order structures ^35,37,100^, although the mechanism underlying stress-induced VIPP1/IM30 puncta and their biological significance remain unclear ^38,101^. We observed prominent VIPP1/IM30 puncta distributed throughout the chloroplast during early stages of thylakoid biogenesis (Fig. S6), suggesting a potential link to membrane remodeling processes. Emerging models propose that VIPP1/IM30 can undergo stimulus-dependent phase transitions, forming liquid-like condensates in response to changes in physicochemical conditions such as pH, ionic strength, and protein concentration ^102,103^. In this context, we propose that VIPP1/IM30 proteins are initially present in a more diffuse, polymeric state at 0 h, and that during early greening, the changes in the local physicochemical environment promote its reorganization into punctate condensates. Such a transition may facilitate membrane remodeling and NT fusion, consistent with recent mechanistic hypotheses ^102,104,105^.

In addition to membrane development, our time-resolved data also elucidate the stepwise assembly of photosynthetic complexes and associated factors, enabling us to delineate the temporal pathways of PSII and PSI supercomplex assembly (Fig. 9b). During the initial stage of thylakoid biogenesis (0-1 h), subunits constituting the PSII and PSI RCs, including PsaA, PsaB, PsbA, and PsbD–F, are synthesized. CP47 and other peripheral PSII subunits rapidly associate with the PSII RC to form a stable RC47 intermediate after 2 h. Subsequently, incorporation of CP43 and the pre-assembled OEC leads to the formation of the PSII core, accompanied by a marked increase in oxygen evolution around 4 h. The assembly of PSI complexes appears to lag slightly behind that of PSII (Figs. 7, 8). Between 4-6 h, the PsaC, PsaD, and PsaN subunits assemble into the intermediate PSI* complex ^106^. Concurrently, the LHCB and LHCA levels increase steadily throughout thylakoid development. After 6 h, LHCB associates extensively with the PSII core to generate the PSII-SC, followed by LHCA binding to PSI after ∼8 h to form the PSI-SC.

Our results also reveal that the pyrenoid and its tubules exhibit a high degree of structural integrity, even when the thylakoid membranes surrounding the pyrenoid were substantially diminished (Figs. 3a, 4d). Consistently, proteomic analysis showed that 70% of RuBisCO, 60% of RBMP1/2 and 40% of EPYC1 were retained after dark treatment, along with high levels of enzymes in the Calvin-Benson cycle (Fig. S11b). During greening, the thylakoid membranes around pyrenoid reorganize into ordered structures and re-establish connections with the tubules (Fig. S3). A recent study documented that the tubule structure could be affected by high temperature ^107^. These results suggest that the maintenance and regulation of the pyrenoid and tubules are distinct from those of typical thylakoid membranes outside the pyrenoid.

Some questions remain about the origin and earliest maturation steps of NTs. Our data cannot definitively distinguish pre-existing residual membranes from *de novo* membrane synthesis, nor fully resolve the temporal order of lipid synthesis, morphology development, local translation, and protein assembly at individual sites. Targeted temporal-tracing experiments, such as time-resolved high-resolution imaging, photoconvertible tags to follow assembly factors, and rapid perturbation assays, would directly test these possibilities and clarify causality during thylakoid biogenesis.

In summary, our work provides a quantitative, spatiotemporal framework for globally distributed, multipolar thylakoid membrane biogenesis during greening in the *Chlamydomonas* Δ*chlL* strain. The findings refine our understanding of thylakoid biogenesis and highlight a more dynamic and decentralized biogenesis process that is tightly coordinated with environmental cues and the underlying cellular architecture. Our system is also expected to support a more detailed comprehensive characterization of the stepwise assembly of the photosystems. The conceptual framework established here provides a foundation for dissecting the underlying molecular mechanisms, regulatory principles, and evolution of photosynthetic membrane biogenesis, with potential implications for cell and membrane engineering, synthetic biology, and crop improvement.

## Materials and Methods

See Supplementary Information

## Acknowledgments

We thank the Centre for Proteome Research (CPR) for support with mass spectrometry, the Centre for Cell Imaging (CCI) for support with fluorescence microscopy, the Biomedical Electron Microscopy Unit for support with transmission electron microscopy. We would like to acknowledge the support of the Materials Innovation Factory (MIF) at the University of Liverpool for 77K fluorescence spectroscopy, which is created as part of the UK Research Partnership Investment Fund (UKRPIF) initiative, managed by UKRI Research England. We are grateful to Dr. Daniel Canniffe for assistance with HPLC, and to Tanda Qi for providing the code used in heatmap generation. We acknowledge the Oxford Particle Imaging Centre (OPIC) for access Aquilos 2 and the cryo-EM instrument Krios. We thank L. Carrique, H. Duyvesteyn, and J. Gilchrist for their support in cryo-ET data collection. We acknowledge Diamond Light Source for access to and support of the cryo-EM facilities at the UK national electron Bio-Imaging Centre (eBIC), proposal NT29812.

## Author contributions

L.-N.L., P.Z. and P.J.N. conceived the project. X.G., Z.H., G.N.J., P.J.N., P.Z. and L.-N.L. designed the experiments and analysed the data. X.G. performed most of the molecular, biochemical analysis, spectroscopic and fluorescence microscopy work. Z.H., J.Z., and M.D. conducted the cryo-FIB/SEM volume imaging. Z.H. and J.H. analysed the cryo-FIB/SEM volume imaging data. Z.H. performed cryo-FIB milling and cryo-ET. Z.H. performed the template matching and subtomogram averaging. Z.H. performed the segmentation and analysed the cryo-ET data with help from L.C. M.Q. performed chloroplast genome transformation and mutant construction. Measurements of P700 were conducted by X.G., P.M. and G.N.J.. Low-temperature fluorescence spectroscopy was performed by X.G., K.P. and Y.Z.. Mass spectrometry data collection and analysis were conducted by J.S. and X.G.. Transmission electron microscopy was performed by X.G. and G.F.D.. X.G. and Z.H. prepared figures. X.G., Z.H., P.Z., and L.-N.L. wrote the paper with the contribution of all other authors.

## Data and materials availability

All data are available in the main text or the Supplementary Information. The subtomogram averaged maps are deposited in the public database EMDB under the accession codes: EMD-55458 (Δ*chlL* translating 70S ribosome in dark), EMD-55459 (Δ*chlL* non-translating 70S ribosome in dark), EMD-55461 (Δ*chlL* translating 70S ribosome in light), EMD-55462 (Δ*chlL* non-translating 70S ribosome in light), EMD-55363 (WT translating 70S ribosome in dark), EMD-55464 (Δ*chlL* F-ATPase), EMD-55465 (Δ*chlL* cytoplasmic 80S ribosome), and EMD-55466 (Δ*chlL* RuBisCO), respectively. Mass spectrometry data used for Fig. 7 and Fig. S11 were deposited to the ProteomeXchange Consortium via PRIDE partner repository with the Project accession PXD070305 (https://www.ebi.ac.uk/pride/archive/projects/PXD070305).

## Supplementary Information

Materials and Methods

Figs. S1 to S13

Tables S1 to S4

Movies S1 to S4

