## Supplemental methods and figures for "Spatiotemporal biogenesis of thylakoid membranes in the green alga *Chlamydomonas reinhardtii*"

### Supplementary Information

**The PDF file includes:**

Materials and Methods
Figs. S1 to S13
Tables S1 to S4
Supplementary References

#### Materials and Methods

##### Strains and culture conditions

*Chlamydomonas reinhardtii* CC-1690 wild-type mt$^+$ (Sager 21 gr) strain was obtained from the Chlamydomonas Resource Center (University of Minnesota). For dark treatment, cells were maintained on Tris-acetate-phosphate (TAP) agar 1.5% (w/v) plate under dark for more than four rounds with antibiotics final concentrations: spectinomycin 200 mg·mL$^{-1}$, paromomycin 20 mg·mL$^{-1}$, then growing in TAP liquid medium for 2 days before 20 µmol photons m$^{-2}$ s$^{-1}$ low light treatment. Specially, the dark treated samples were collected under dark and covered by foil during processing.

The Δ*chlL* strain was generated by replacing *chlL* gene on chloroplast genome by a spectinomycin resistance gene (*aadA*). Briefly, a pGEM-T plasmid containing *aadA* coding sequence along with the flanking sequences of *chlL* gene was transferred to CC-1690 chloroplast via biolistic bombardment, following the protocol outlined by [1]. 100 µL-200 µL of the concentrated cells (2 × 10$^7$ cells/ml) were spread on selection plate (TAP with 150 µg·mL$^{-1}$ spectinomycin) and plasmid DNA were delivered into the cells via golden particles using Biolistic PDS-1000/He (BioRad) device. After transformation, plates were incubated overnight under dim light (~ 5 µmol photons m$^{-2}$ s$^{-1}$) and then transferred to moderate light (~ 20 µmol photons m$^{-2}$ s$^{-1}$) the next day. Successful transformants were picked and subjected to several rounds of re-streaking on selection plates for fully segregation. Genotypes of the transformants were confirmed by genotyping (Fig. S1) and sequencing.

Genomic DNA was extracted from *C. reinhardtii* using phenol-chloroform. For fluorescent labelling strains, full-length genomic sequences of the targeted genes, containing native exons, introns, and transit peptides (ensuring correct chloroplast transport), were amplified from purified CC-1690 genome by PCR using CloneAmp HiFi (Takara) and were inserted in-frame with a C-terminal Venus into the pLM005 plasmid by Gibson assembly, driven by the strong PsaD promoter [2-5]. A flexible linker peptide (GDLGGSGG) was introduced between the target protein and Venus to minimize steric hindrance and facilitate proper protein folding. For EPYC1-crVenus labelling, the pLM017 plasmid that derived from pLM005 was used as previously reported [2]. DNA fragments containing crVenus-fused cassette and paromomycin resistance gene (controlled by the RBCS2 promoter) were amplified by PCR and were transformed to Δ*chlL* by nuclear transformation. In short, Δ*chlL* cells were grown under light and collected by centrifugation at 2,000 × g, followed by washing with fresh TAP medium with 50mM sucrose, 1 µg of DNA was mixed with 10$^8$ cells, electroporated with Bio-Rad Gene Pulser Electroporation system (Bio-Rad) with 800 V, 25 µF, 800 Ω. Colonies appeared after 5 days and positive transformants were finally confirmed by detection of crVenus signals using LSM 780 (ZEISS) with 516 nm laser.

**Pigment qualification**
Liquid cultures were collected, and pigments were extracted by HPLC grade methanol, the supernatant of extraction were analysed by Cary UV-Vis compact peltier (Agilent) at 470, 652, 665 nm. Contents of Chl *a*, Chl *b*, and carotenoid were calculated as reported [6] with following equations: Chl $a = 16.29 \times A_{665} - 8.54 \times A_{652}$, and Chl $b = 30.66 \text{ x } A_{652} - 13.58 \times A_{665}$, and carotenoids = $(1000 \times A_{470} - 2.86 \times \text{Chl } a - 129.2 \times \text{Chl } b)/221$. For HPLC, Chl was extracted from equal cells (same $OD_{750}$) with HPLC grade methanol (0.2% ammonium hydroxide) and analysed by C18 Reversed Phase Columns.

**Protein gel and immunoblot analysis**
Membranes were isolated from Δ*chlL* cells treated with low light using method reported [7]. Proteins were released from membranes by 30min 1% n-dodecyl-β-D-maltoside treatment in final concentration. Insoluble materials were removed by 18,000 × *g* for 20 min centrifuge at 4°C. The supernatant was mixed with 10X blue native loading buffer and 150 µg proteins were load to 4–15% Mini-PROTEAN Protein Gels (Bio-Rad) for analysis, running voltage was gradually increased from 50 V up to 150 V. The bule cathode buffer was replaced by clear cathode buffer without Coomassie Blue when the dye bands went to half of the gel. The proteins were transferred to PVDF membranes (Amersham™) for immunoblotting analysis with primary antibody α-PsbA (AS05 084, Agrisera) in dilution of 1:5000, α-PsaA (AS06 172, Agrisera) in dilution of 1:5000 and secondary antibody (AS09 602, Goat anti-Rabbit, Agrisera) in dilution of 1:10000. ImageQuant LAS 4000 v1.2.1.119 was used to record chemiluminescence signal by ECL (Bio-Rad).

**Transmission electron microscopy**
For thin section transmission electron microscopy (TEM) analysis, cells were harvested from 100 mL liquid cultures with light-shaded operation. Cells were fixed with 2% glutaraldehyde in 0.1 M sodium cacodylate buffer immediately and stained by 2% OsO4 (Osmium) 1% TCH (Thiocarbohydrazide) and by 1% UA (Uranyl Acetate) overnight. Cells were dehydrated with alcohol and acetone, fixed in medium resin (TAAB) for polymerization. 70nm thin sections produced through diamond knife were collected by copper grids and post-stained with 3% lead citrate. Images were collected using a FEI Tecnai G2 Spirit BioTWIN transmission electron microscope equipped with a Gatan Rio 16 camera under 120 kV.

**Plunge-freezing vitrification**
An aliquot of 3.5 µL *C. reinhardtii* cell suspension at $1.5 \text{x} 10^6$ cells/ml was applied to the glow-discharged holey carbon-coated copper grid (R 2/1, 200 mesh) (Quantifoil) and blotted on the back of the grid for 9 seconds by Leica GP2 (Leica Microsystems), followed by the plunge freezing in liquid ethane.

**Cryo-FIB/SEM volume imaging**
The cryo-FIB/SEM volume imaging was performed on a dual-beam FIB/SEM microscope Helios G4 Hydra equipped with a cryo-stage cooled at -191℃ by an open nitrogen circuit, and a plasma multi-ion source (argon, nitrogen, xenon, and oxygen) Hydra plasma ion column and an

Elstar SEM column (Thermo Fisher Scientific) (Table S1). An immersion field is generated to improve detection and signal to noise. The samples were loaded into a custom purpose 25° pre-tilt holder. To reduce the charging during SEM acquisition, sputtering coating was performed for 50 seconds before and after the deposition of an organic platinum layer. The deposition of organic platinum layer was performed by the gas injection system (GIS) for 50 seconds to protect the front of samples from beam damaging. Cells positioned in the centres of grid squares were selected and the opening of the cells was conducted by FIB at 30 kV and 2 nA using argon as source, removing the top materials, followed by a polishing at 30 kV and 200 pA. To suppress the drift during the acquisition, a fiducial mark was produced on one side of the sample. Then the milling and imaging sequence commenced, in which the FIB milling was performed using argon at 30 kV and 30 pA, removing 20 nm in thickness of the sample per cut, the SEM imaging was set at 1.0 kV and 6.3 pA with 100-line integration at 4.5 nm/pixel. The process was automated using Auto Slice and View (ASV) version 5.0 (Thermo Fisher Scientific), summarised in (Table S1).

**Cryo-focused ion beam milling for lamellae production**

Vitrified cells were further thinned by cryo-FIB milling for the preparation of lamellae. WT cells were loaded and prepared on a dual-beam FIB/SEM microscope equipped with a cryogenic stage cooled at -191℃, fluorescence microscope, and a plasma multi-ion source (argon, nitrogen, xenon, and oxygen), which is the prototype of commercially available microscope Arctis Thermo Fisher Scientific), argon was used as the FIB source. *ΔchlL* cells were thinned by a dual-beam FIB/SEM microscope Aquilos 2 (Thermo Fisher Scientific) equipped with cryo-transfer system (Thermo Fisher Scientific) and rotatable cryo-stage cooled at -191℃ by an open nitrogen circuit. Prior to the milling, the grids were mounted on the shuttle and transferred onto the cryo-stage, followed by the coating with an organometallic Platinum layer using the GIS system (Thermo Fisher Scientific) for 20 seconds. Then, cells positioned approximately in centres of grid squares were selected for the lamellae production. The thinning was conducted by the automated milling software AutoTEM 5 (Thermo Fisher Scientific) in a stepwise manner from current 0.5 nA to 30 pA at 30 kV, and the final thickness of lamellae was set to 120 nm, summarized in (Table S1).

**Cryo-electron tomography**

Cellular lamellae were transferred to a FEI Titan Krios G3 (Thermo Fisher Scientific) electron microscope operated at 300 kV and equipped with a Falcon 4i detector and a Selectris X energy filter (Thermo Fisher Scientific). A 100 µm objective aperture was inserted. Overviews of each cellular lamella was acquired, and chloroplast regions were targeted for the following tomography. For the WT, tilt series were collected using Tomography 5 software (Thermo Fisher Scientific) with a nominal magnification of 64k and a physical pixel size of 1.978 Å/pixel. For the *ΔchlL* in dark and after 1-hour light treatment, tilt series were collected using Tomography 5 software (Thermo Fisher Scientific) with a nominal magnification of 64k and a physical pixel size of 1.94 Å/pixel. For the *ΔchlL* after 24-hour light treatment, tilt series were collected using Tomography 5 software (Thermo Fisher Scientific) with a nominal magnification of 64k and a physical pixel size of 1.9026 Å/pixel. All tilt series were collected with a zero-loss imaging filter with a 10 eV-wide slit. The defocus value was set from -3 to -5 µm. The pre-tilt of the lamellae was determined at ± 9°, and a dose-symmetric scheme was applied for all tilt series, ranging from -45° to +63° or -63° to +45°with an increment of 3°. A total of 37 projection images with 10 movie frames each were collected for each tilt series, and the dose rate was set 1.75 e/Å2/s with the exposure time of 2 seconds, resulting in a total dose of 129.5 e/Å2. For the WT, a total of 118 tilt series were collected from 39 lamellae. For dark-treated *ΔchlL* cells, a total of 152 tilt series were collected from 17 lamellae. For *ΔchlL* cells with 1-hour light treatment, a total of 163 tilt series were collected from 21 lamellae. For *ΔchlL* cells with 24-hour light treatment, a total of 31 tilt series were collected from 8 lamellae. Data collection is summarised in (Table S2).

**Alignment of cryo-FIB/SEM volume images**
Acquired volume images were pre-processed in ImageJ adjusting brightness and contrast, the mean filter of 1 pixel was applied. Pre-processed images were then imported into IMOD [8] version 4.11.1 and aligned using the function Align Serial Sections / Blend Montages. The Initial auto alignment was performed followed by the refinement with Midas, and the final stack was created with all trending removed.

**Alignment of tilt series and tomogram reconstruction**
The frames of each tilt series were corrected for beam-induced motion using MotionCor2 [9]. New stacks were generated and aligned in IMOD version 4.11.1 by patch tracking using patches of 200 × 200 pixels and a fractional overlap of 0.45 in X and Y, and tomograms were reconstructed at bin6 with a pixel size of 11.868 Å/pixel for the WT and 11.64/11.4146 Å/pixel for *ΔchlL*. For a better visualization, SIRT-like filtering was applied to reconstructed tomograms with 8 iterations.

**Template matching**
To localize chloroplast 70S ribosomes, F-ATPase, cytoplasmic 80S ribosomes, and RuBisCO particles in the tomogram, template matching was carried out using emClarity [10] version 1.5.0.2 on well-aligned tomograms with low residual error weighted means (< 0.8 nm) in the alignment. To supress the template-induced bias, a low-pass filtering at 40 Å was applied to the templates: EMD-1780 [11] (80S ribosome), EMD-3533 [12] (70S ribosome), and EMD-22401 [13] (RuBisCO). For 70S ribosomes, 142 tomograms were selected for dark-treated *ΔchlL* cells, 64 tomograms were selected for light-treated *ΔchlL* cells, and 28 tomograms were selected for dark-treated WT cells. Initially, 22,800, 9,600, and 4,200 particles were picked for these three conditions, respectively. For F-ATPase, to mitigate template-induced bias, a featureless sphere approximating the size of F-ATPase head was employed as the initial template (Fig. S10a) in 32 tomograms, yielding 12,800 particles initially. For 80S ribosomes, total number of 10 tomograms were selected for template matching, yielding 4,000 particles in total after cleaning by manual inspection. For RuBisCO, 40 tomograms were selected, and 397,481 particles were extracted after manual cleaning.

**Subtomogram averaging**
Before aligning the particles, CTF correction was performed for each tomogram by emClarity version 1.5.3.10, and particles were inspected by overlaying the reconstructed tomograms with corresponding picked particles in Chimera for further cleaning. Particles residing in non-corresponding cellular compartments were removed. Remaining particles were then imported into RELION [14] version 4.0 for iterative reconstructions and alignments from binning at 6 to binning at 2. For each macromolecule, 3D classification was conducted at the binning of 6 to investigate the heterogeneity. For 70S ribosomes, a mask was generated covering the P site for the focused classification (Fig. S9a-c) to distinguish the translation states, the structure of dark-

treated *ΔchlL* non-translating 70S ribosome was resolved at 6.4 Å with 9,757 particles, the translating one was resolved at 10.0 Å with 2,501 particles; the structure of light-treated *ΔchlL* non-translating 70S ribosome was resolved at 16.9 Å with 819 particles, the translating one was resolved at 13.3 Å with 3,992 particles; the structure of dark-treated WT translating 70S ribosome was resolved at 19.5 Å with 1,653 particles (Fig. S9d). For F-ATPase, 2,026 particles with prominent density of the stem connected with bilipid layer were selected after 3D classification (Fig. S10a, b), the structure was determined at 24.8 Å (Fig. S10f). As the subunit d was not fully incorporated into the complex in dark-treated *ΔchlL* cells and *ΔchlL* cells with 1-hour light treatment (Fig. S10 b), leading to the instability of the overall structure, the flexibility of the final structure was high. For RuBisCO particles, the heterogeneity was high, and the structure was determined at 8.5 Å (Fig. S10 d, f) with 21,560 particles after several rounds of cleaning through 3D classification. For 80S ribosomes, after 3D classification, 2,950 particles were included for the final structural determination, and the resolution reached 10.6 Å (gold-standard 0.143 cut-off) (Fig. S10 e, f). Information is also summarised in Table S3.

**Segmentation of tomograms**

To enhance the segmentation, reconstructed tomograms were corrected for missing wedge and denoised by IsoNet [15] version 0.2, applying 35 iterations with a sequential noise cut-off level of 0.05, 0.1, 0.15, 0.2, 0.25 at iteration 10, 15, 20, 25, 30. For the WT, three tomograms with typical thylakoid stacks were selected. For dark-treated *ΔchlL* cells, *ΔchlL* cells with 1-hour and 24-hour light treatment, nine, seven, and three tomograms covering different regions (basal region, lobe, T zone, and pyrenoid) were selected, respectively. Membranes in all tomograms were initially segmented using MemBrain-seg [16] and then imported into ChimeraX [17] for cleaning and polishing. 80S ribosomes, 70S ribosomes, RuBisCO particles, and F-ATPases were mapped back to the tomograms with segmented membranes using ChimeraX and ArtiaX [18] based on their refined positions and orientations. For a better visualization, the macromolecule models depicted in the segmented volume were generated by applying a low-pass filter with a cut-off of 15 Å to their respective templates.

**Measurement of thylakoid curvature, intermembrane distance, and lumenal density**

To measure the overall curvature of thylakoid membranes, nine tomograms of dark-treated *ΔchlL* cells, seven tomograms of *ΔchlL* cells with 1-hour light treatment, three tomograms of *ΔchlL* cells with 24-hour light treatment, and three tomograms of WT cells were segmented and polished using the same parameter as described in the segmentation section. Then only thylakoid membranes were kept for the measurement of mean curvature using Pyvista [19] version 0.43 (https://docs.pyvista.org/version/stable/). As the edges of segmented thylakoid membranes near the boundary of tomograms were highly curved, induced by the segmentation per se, edges were then excluded from the measurement. In total, 1,665,207 sampling points from the dark-treated *ΔchlL* cells, 4,890,773 sampling points from the 1-hour-light-treated *ΔchlL* cells, 7,556,984 sampling points from the 24-hour-light-treated *ΔchlL* cells, and 8,679,442 sampling points from the WT cell were extracted. The mean curvature of each sampling point was then calculated. To measure the intermembrane distance of thylakoid, thylakoid fragments were randomly selected from reconstructed tomograms (bin6) of the four samples. The measurement of intermembrane distance commenced by manually marking points along one of the paired membranes of selected thylakoid fragments. These marked points served as a basis for curve fitting. Polynomial functions were subsequently applied to fit curves independently for each membrane. Following the curve fitting, points on the membrane were sampled, and normal lines were generated at these points, intersecting with the curve of the other membrane. Distances between each sampled point and its corresponding intersection point on the other membrane were then calculated. This entire procedure was repeated for the other membrane of selected thylakoid fragments to ensure the intermembrane distance was measured in both directions. In total, 320 paired points were generated for each sample. The intermembrane distance was ultimately determined by subtracting the mean thickness of the lipid bilayer (approximately 5.5 nm) from the point-to-point distance. For both statistical analyses, non-parametric ANOVA (Kruskal-Wallis) test was applied to investigate the statistical difference between the four samples, and the box plot was used to for the illustration.

To quantify the lumenal density of thylakoids across different stages of the thylakoid biogenesis. Twenty thylakoid membranes from each sample were selected and the lumenal areas were framed, followed by the intensity measurement in ImageJ [20]. Statistical plots were performed using Prism [21] version 10 (https://www.graphpad.com/updates/prism-1000-release-notes).

**Calculation of particle densities on the membrane**

Cleaned segmented membranes were read into MATLAB version R2022R, and the voxels of segmented volumes (value read > 0) were calculated. The volume of membranes was calculated by multiplying the voxel number by the voxel size (11.64 × 11.64 × 11.64 Å). Then the surface area was calculated by dividing the volume by the measured mean thickness of lipid bilayer (approximately 5.5 nm) and then multiplying the result by factor of 2. At last, particles on the membrane were counted, and the density was obtained by dividing the particle number by half of the measured membrane area as there was only one surface embedded with F-ATPases.

**Super-resolution fluorescence microscopy**

Cells were pipetted on top of TAP agar plate under dark for immobilization, then an agar square with dropped cells was cut out and placed against coverslip for imaging by Elyra 7 system (ZEISS) equipped with 63×/1.4 NA oil immersion objective. 13 phrase images were taken for each snap under Lattice model with 1280 X 1280 pixel 16 bits. To prevent signal leaking and improve signal collection, SBS LP560 filter was applied to assign wavelength for dual-camera system with laser 488nm for crVenus captured by TV2 and laser 642nm for Chl fluorescence captured by TV1. Dual-camera alignment was performed with standard pattern ahead imaging session. Finally, by ZEN Black software, $SIM^2$ deconvolution reconstruction was used to generate super-resolution fluorescence images with standard parameters: iterations 16, regularization weight 0.0650.

**Whole cell absorption spectrum measurement and 77 K fluorescence spectra**

Whole-cell absorption spectra measurement was carried in a 1cm cuvette at room temperature using Cary UV-Vis compact peltier (Agilent) with 1 nm increments, blanked with TAP medium. Fluorescence emission/excitation spectrum was recorded at 77 K with an FLS-1000 (Edinburgh Instruments) equipped with a liquid nitrogen cryostat (Oxford Instruments). Cells were resuspended with 60% glycerol and frozen in liquid nitrogen for 15 minutes in a 2 mL cuvette covered and transferred under nitrogen atmosphere to a precooled cryostat. The sample is excited at 440 nm for Chl *a* and 475 nm for Chl *b*. Band width for excitation and emission is set to 1 nm and dwell time set to 0.20s. Emission spectra is collected with 739nm/715nm for PSI, and red-shifting PSII emission peak from each time point.

**PSII activity**
Oxygen evolution activity was analysed by Clarke-type OxyLab2 system (Hansatech) at 25°C. Oxygen evolution rate was calculated from oxygen taken and release cycle from three independent biological replicates and normalized to $OD_{750}$. PSII maximum efficiency Fv/Fm was measured with 1ml culture collected at each time point. Measured using an AquaPen-C fluorometer (Photon Systems Instruments, Brno, Czech Republic) after dark adaptation for three minutes.

**Mass spectroscopy**
Cells were centrifuged, washed with PBS and immediately frozen in liquid nitrogen until use. Cells were resuspended in Lysis buffer (1% SDS, 1% Igepal 1% Sodium Deoxycholate, 125 mM NaCl, 5 mM EDTA, 100 mM Tris-buffered to pH=8.0). They were boiled at 80 ˚C for 10 minutes and then sonicated on ice using a probe sonicator for 5 cycles of 20 seconds on and 10 seconds off at an amplitude of 30%. Protein samples were quantified by BCA Assay. Samples were reduced in 4 mM DTT and alkylated with 14 mM iodoacetamide. Samples were digested with trypsin and purified using SP3 beads [22].

Samples were analyzed using an Evosep One Liquid chromatography system (Evosep Biosystems, Odense, Denmark) coupled online to a TIMS ToF HT mass spectrometer (Bruker) using the inbuilt 30 SPD (samples per day) method. Loading approximately 300 ng of peptides. Mass spectrometry analysis was performed in dia-PASEF mode with a m/z range of 400-1201 and a mobility range of $1/K_0$ = 0.6-1.6 V/cm$^2$ using equal ion accumulation and ramp times of 100 ms. The collision energy was lowered as a function of increasing ion mobility from 59 eV at $1/K_0$ = 1.6 V/cm$^2$ to 20 eV at $1/K_0$ = 0.6 V/cm$^2$. The dia-PASEF window scheme consisted of 32 isolation windows of 26 Da width (estimated cycle time of 1.8 s) with 1 Da mass overlap, and 1 Mobility Window with no overlap.

Data was analysed by Spectronaut (version 19.1, Biognosys), searched against the *C. reinharditt* taxid 3055 (downloaded on the 11Nov2024). The proteins related to thylakoids (Table S4) [23-27] were manually checked refer to Chlamylibrary (https://www.chlamylibrary.org/), genome [28] and chloroplast genome (NC_005353) the protein names were re-annotated according to the Protein Assession in uniport (https://www.uniprot.org/).

**PAM measurement**
P700 signal was recorded by Dual KLAS NIR (Heinz Walz GmbH) using single model, 820-870. The cells were concentrated to approximately $0.5\times10^8$/mL in the ED-101US/MD accessory cuvette using TAP containing 40% sucrose, supplemented with 20 mM NaHCO3 and dark-adapted for at least ten minutes until the baseline was stable. 26000 µE saturating pulse was given 200 ms for each saturation pulse (SP) kinetic measurement to oxidize P700 of PSI, causing absorbance changes in the far-red region. The absorbance difference between 820 nm and 870 nm was used to characterize the oxidation and reduction of P700.

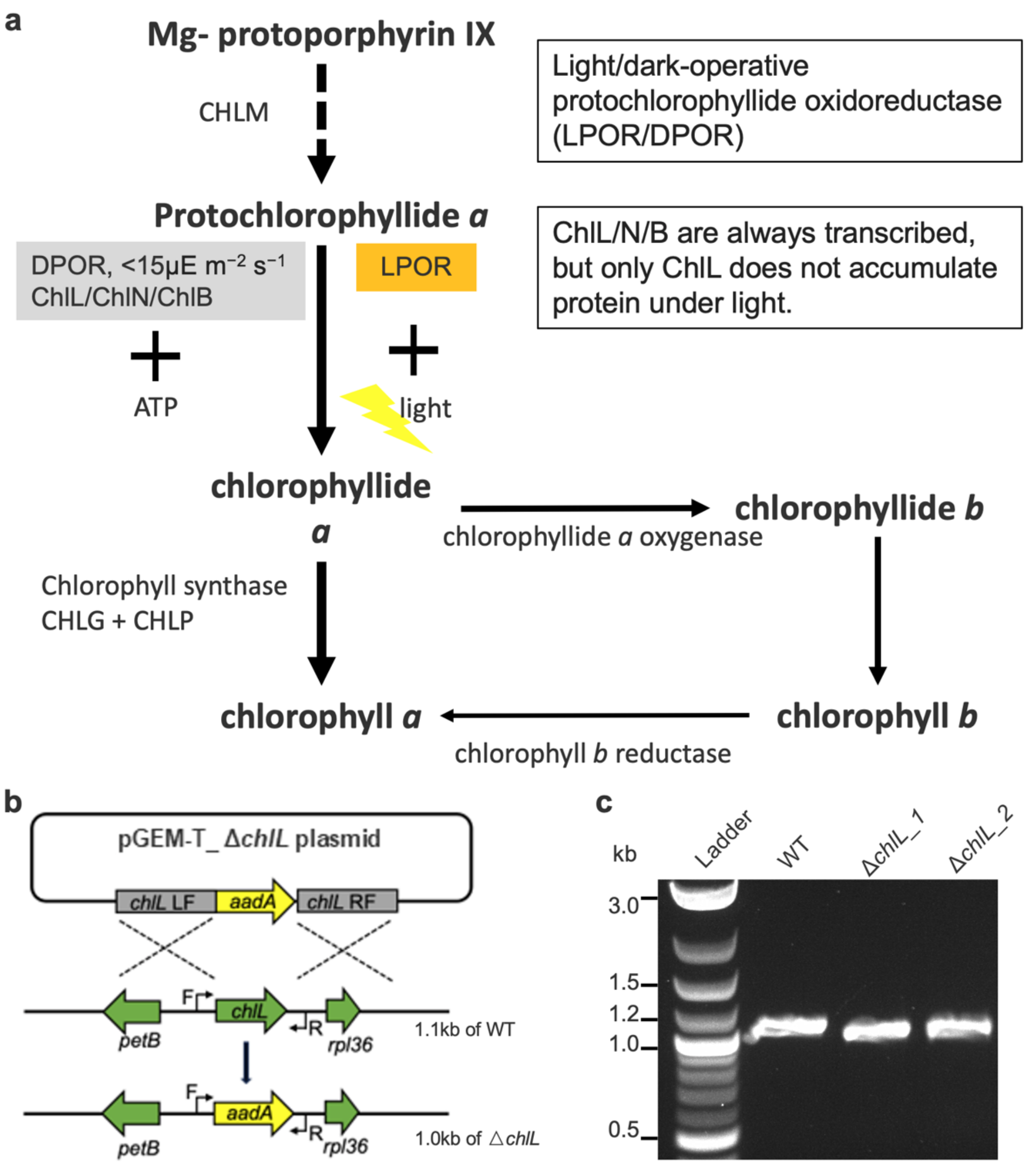

**Fig. S1. Blocking the light-independent Chl synthesis pathway in *C. reinhardtii*. a**, Schematic diagram of the Chl synthesis pathway. The necessary condition for LPOR is light, while DPOR requires three proteins: ChlL, ChlB, and ChlN. **b**, The *chlL*-deletion strategy by homologous recombination. The grey indicates flanking sequences for homologous recombination of *chlL*. Black arrows F and R indicating the binding site of segregation primers. **c**, Genotyping of the Δ*chlL* mutants using PCR and agarose gel electrophoresis.

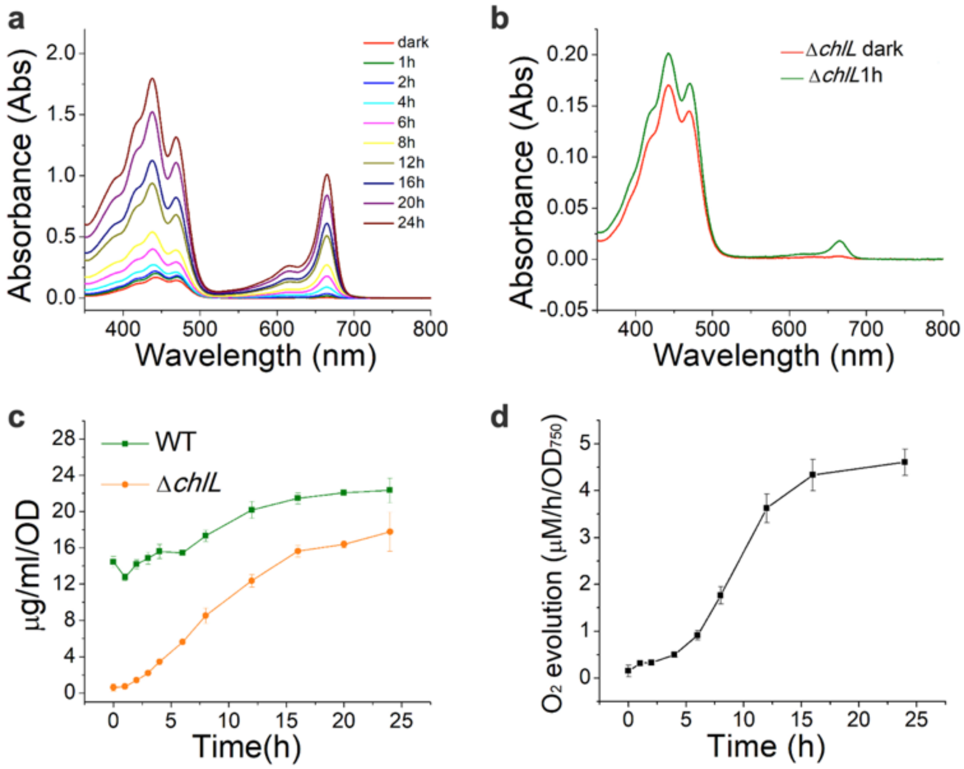

**Fig. S2. Pigment analysis and photosystem activity assays during greening. a-b**, Pigments extraction of Δ*chlL* cells using methanol at different greening timepoints. Room-temperature absorption spectra were recorded with four repeats. All spectra were normalized to 750 nm. **c**, The chlorophyll content was measured by MeOH extraction. Values are means ± SD; n = 3 biologically independent preparations. The content was normalized to $OD_{750}$ to present average chlorophyll content in cells during greening. **d**, Oxygen-evolution rates measured by oxygen2 lab system. The rate was calculated from at least 3 respiration and oxygenation cycle and normalized to cell density.

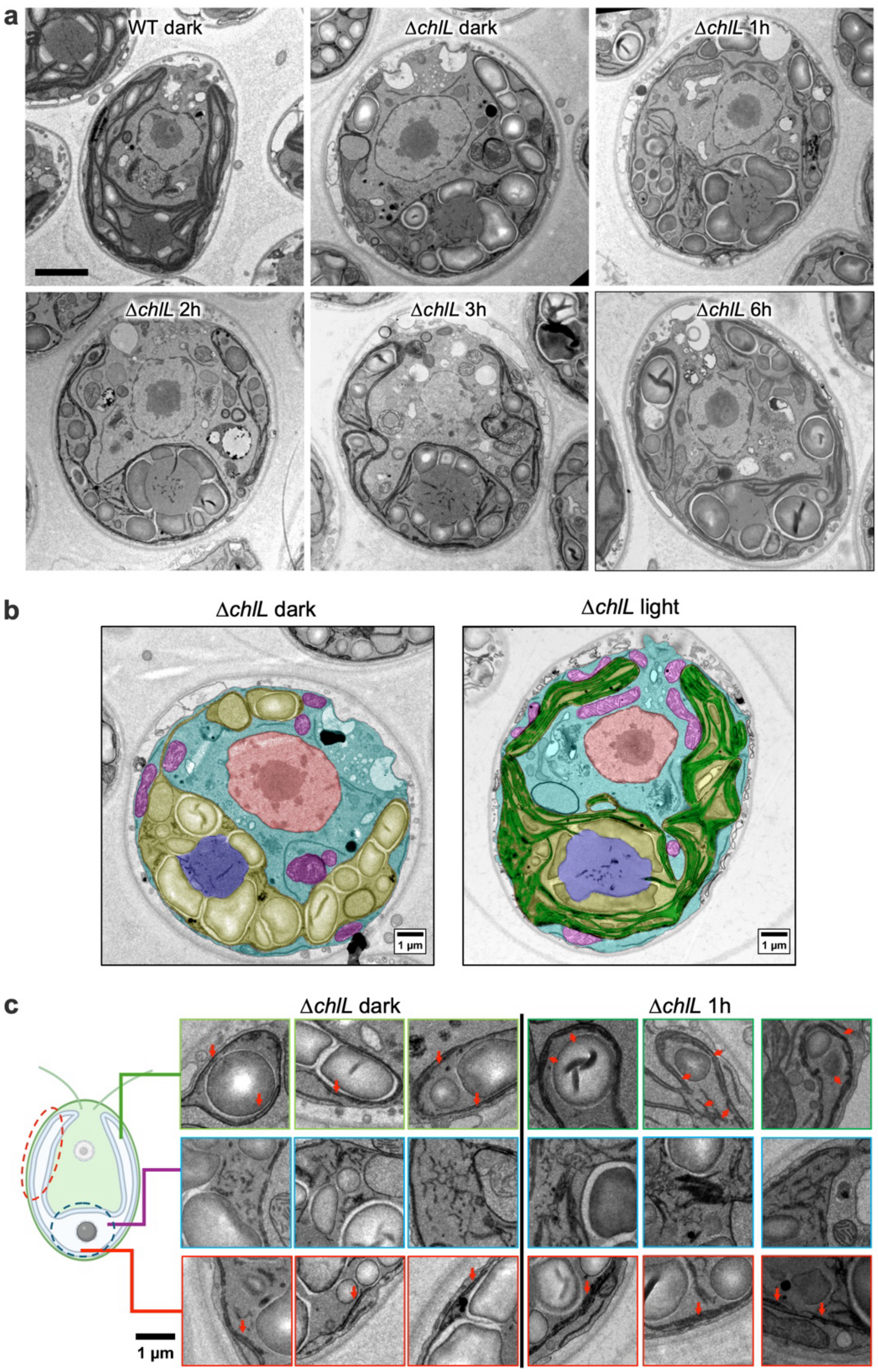

**Fig. S3. Characterization of thylakoid development during greening. a**, Raw thin-section TEM images of *C. reinhardtii* WT and the Δ*chlL* mutant grown under dark or light conditions, related to Fig. 1f. Scale bar = 2 µm. **b**, Representative thin-section TEM images of the Δ*chlL* cells grown under dark or light conditions, with the nucleus coloured red, the mitochondria coloured purple, the cytoplasm coloured cyan, the chloroplasts coloured light yellow, and the pyrenoid coloured blue. **c**, Representative zoomed-in TEM images of the *Chlamydomonas* chloroplast regions, with green for lobe, blue for T-zone stroma, and red for basal. Red arrows indicate traces of thylakoid membranes.

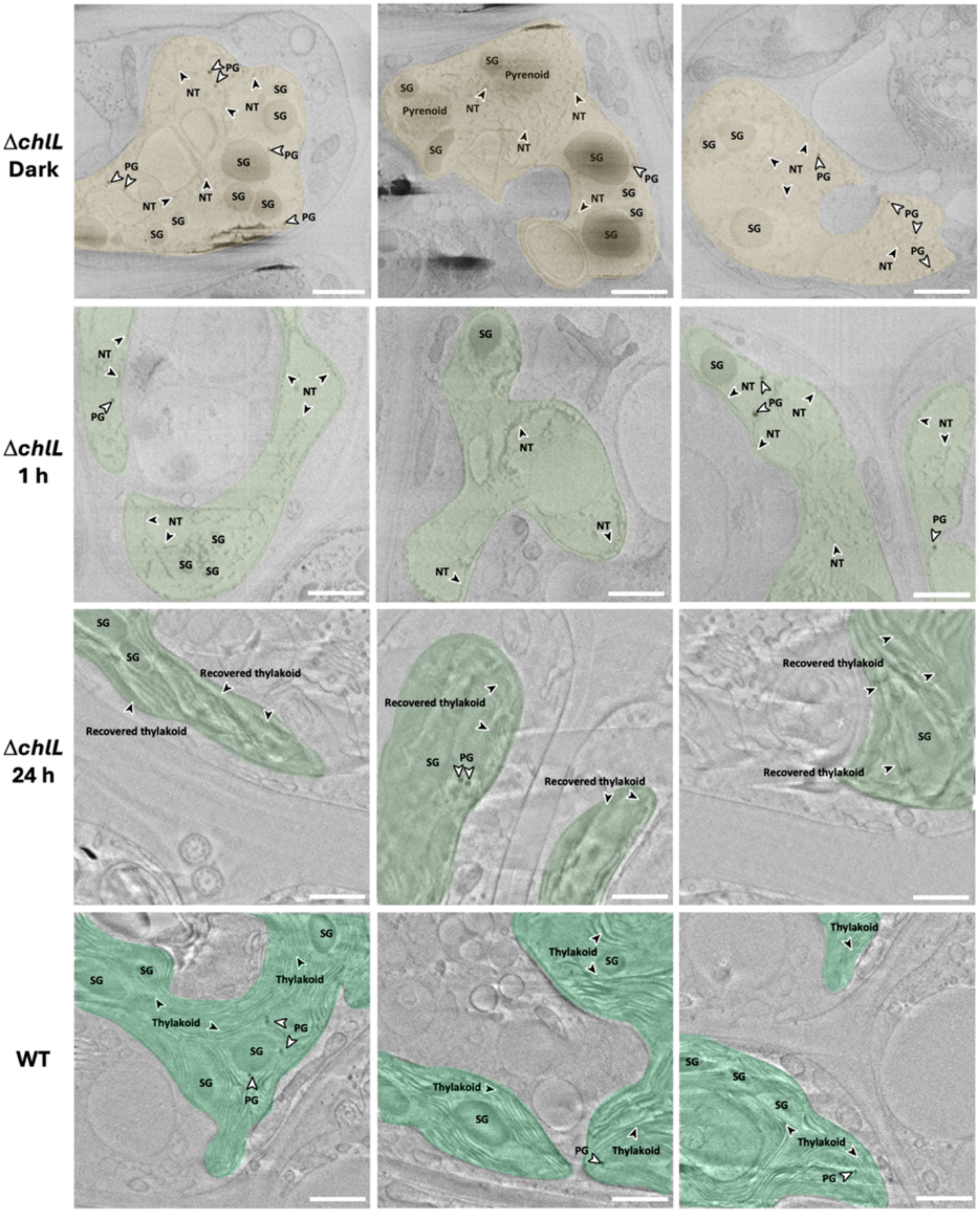

**Fig. S4. Gallery of cryoFIB/SEM volume images of Δ*chlL* and WT.** Representative volume slices of Δ*chlL* dark, Δ*chlL* 1 h and Δ*chlL* 24 h light treatment, and WT cells. The chloroplast region is coloured light yellow or green, and nascent thylakoids (NT) are indicated by black arrowheads, plastoglobules (PG) are indicated by white arrowheads, starch granule: SG. Scale bar = 1 μm.

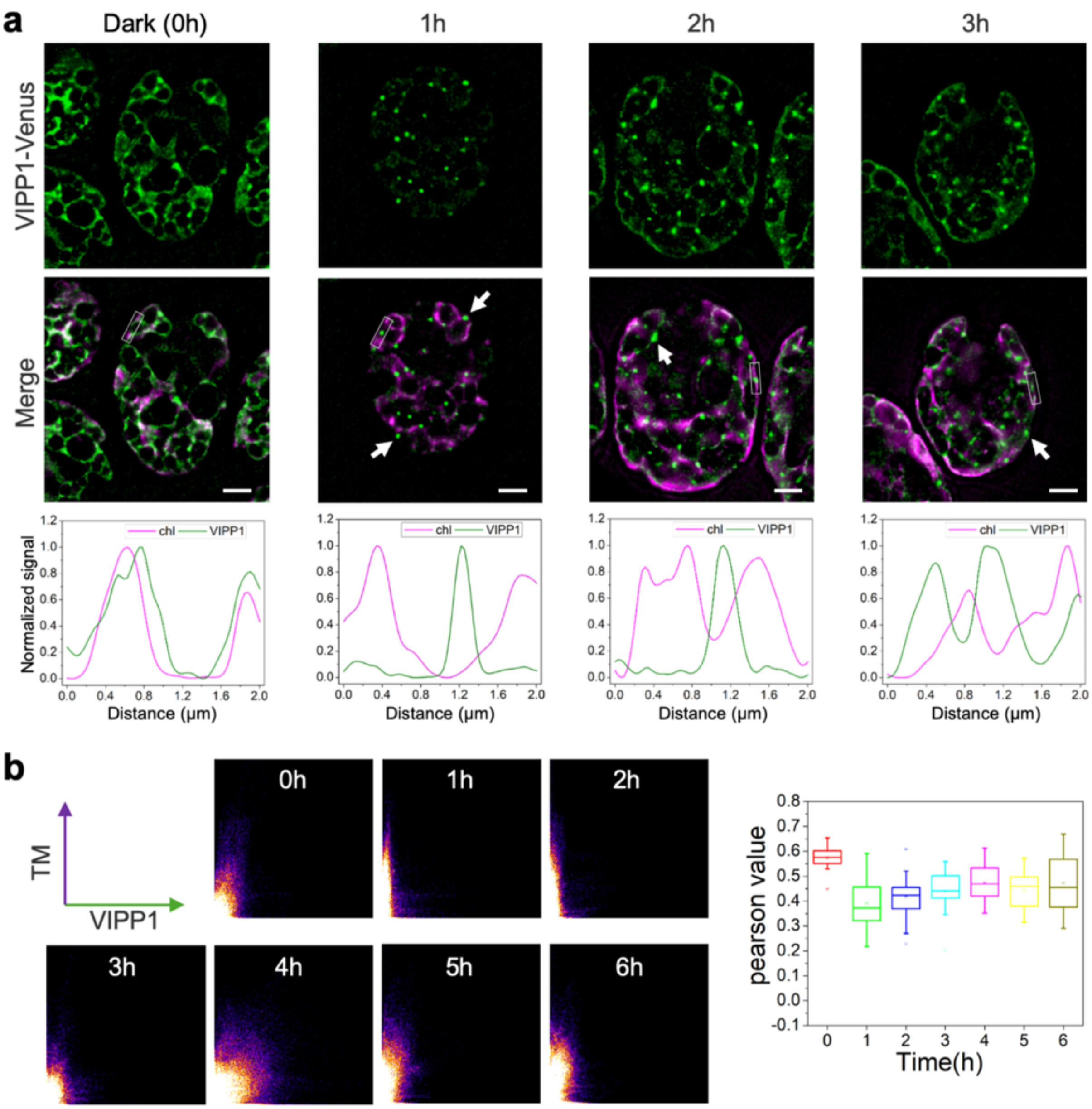

**Fig. S5. Distribution of Vipp1 proteins during thylakoid biogenesis.** **a**, Super-resolution fluorescence imaging of Δ*chlL* VIPP1-Venus during thylakoid biogenesis. The signal intensity at the white box was recorded and shown below the fluorescence images, normalized to the maximum value of 1 to present relative localization of VIPP1 and Chl. White arrows indicate the VIPP1 puncta. Scale bar = 2 μm. **b**, Colocalization analysis of Chl signal and VIPP1 signal, scatterplot was generated from representative images, Pearson's correlation values shown in box plots with the median (line), the average (empty square), the interquartile range (box), and the whiskers (extending 1.5 times the interquartile range).

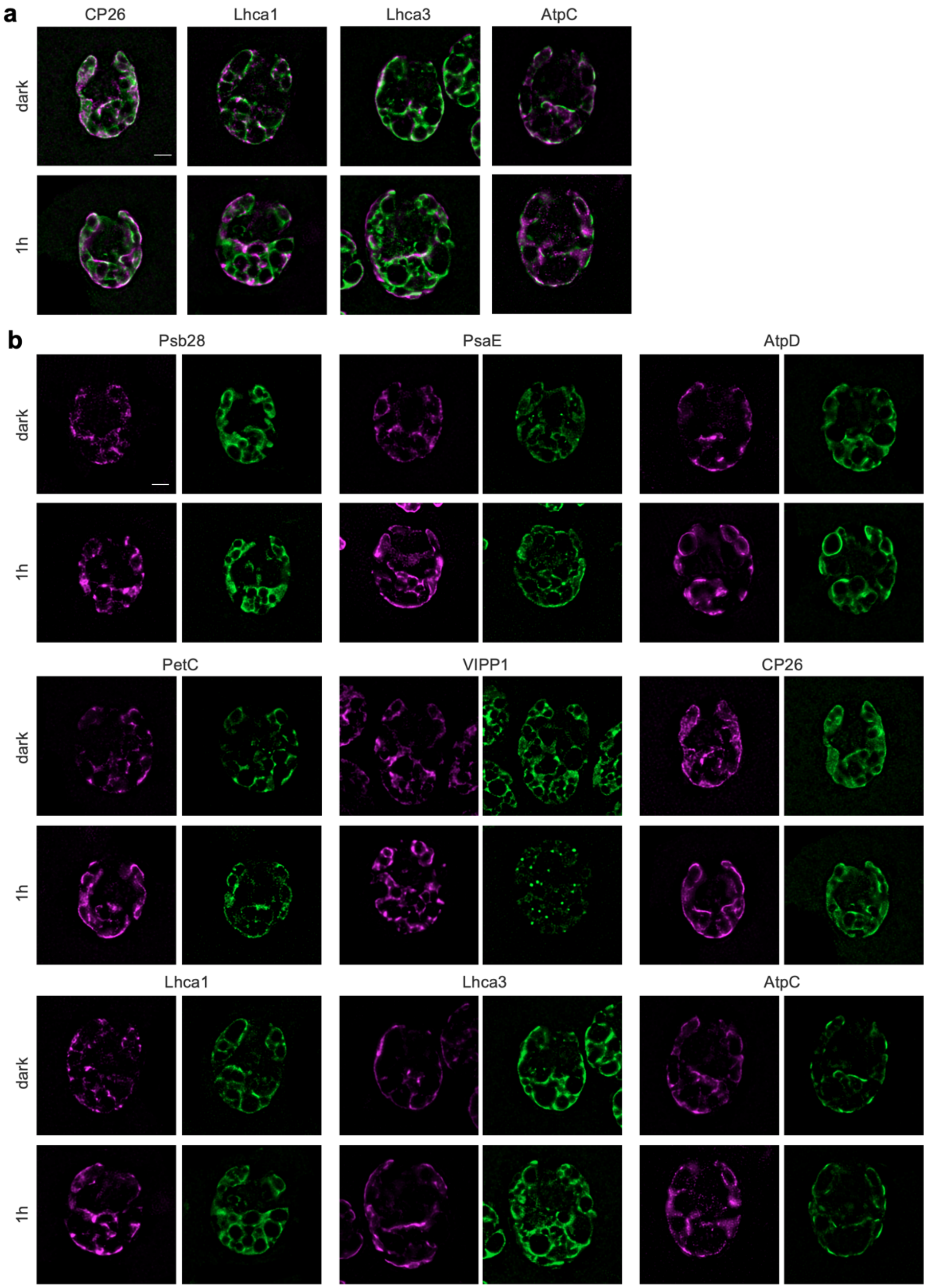

**Fig. S6. Localization of photosynthesis-related proteins during thylakoid biogenesis. a**, Representative merged images of Δ*chlL* CP26-Venus, Δ*chlL* Lhca1-Venus, Δ*chlL* Lhca3-Venus and Δ*chlL* AtpC-Venus, Chl fluorescence coloured in purple, Venus in green, Chl signal brightness was adjusted to illustrate relative localization. Scale bar = 2 µm. **b**, The split-channel images of Fig. 3b and Fig. S6a, the signal intensity was adjusted to saturation to highlight the signal localization in the whole chloroplast. Left panel is Chl fluorescence and the right panel is Venus fluorescence. Scale bar = 2 µm.

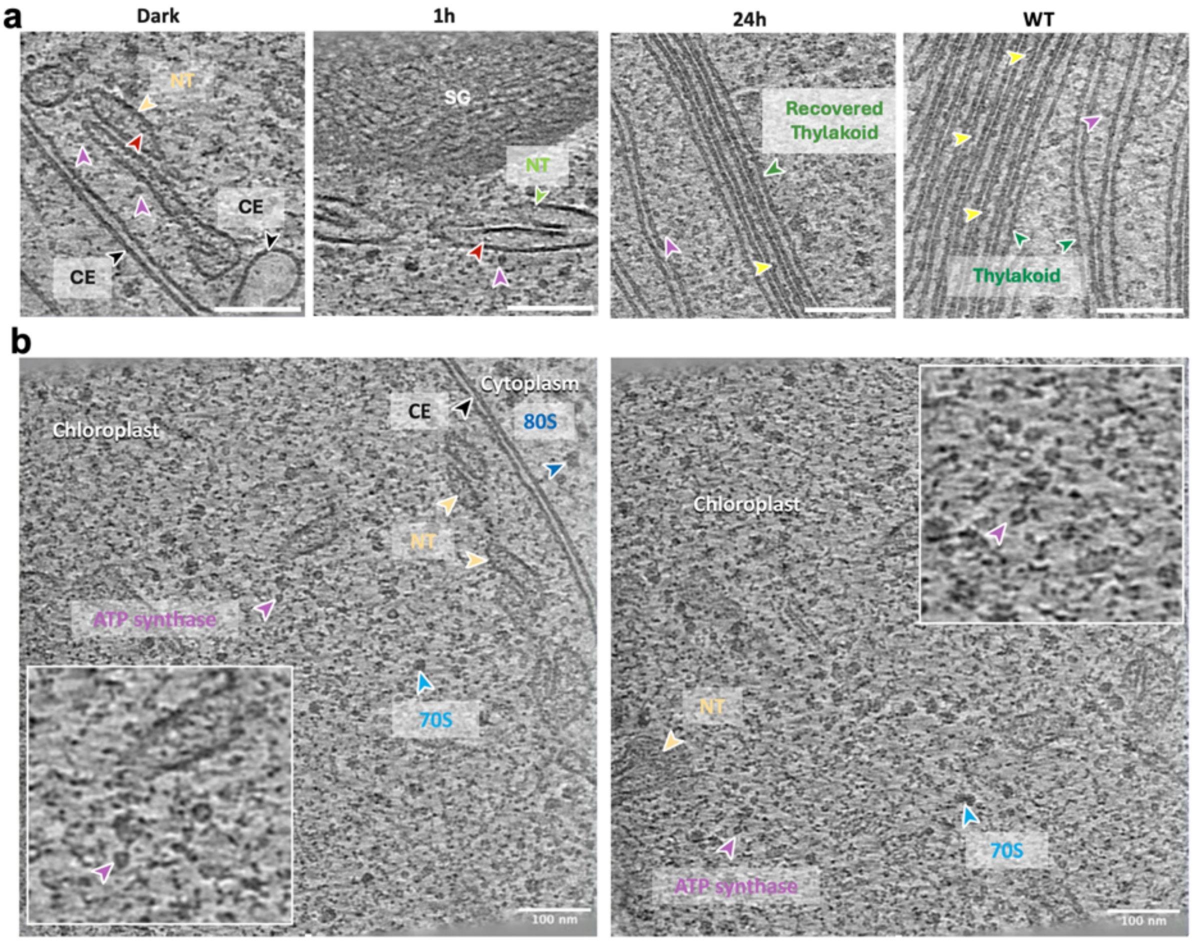

**Fig. S7. Cryo-ET of protein complexes in the Δ*chlL* cells. a**, Zoomed-in tomographic slices featuring the lumenal density and discernible photosynthetic complexes on NTs and mature thylakoids. Lumenal densities are indicated by red arrowheads, F-ATPases are indicated by purple arrowheads, and other photosynthetic complexes are indicated by light yellow arrow heads. **b**, Two representative tomographic slices featuring the membrane-free F-ATPases in the stroma. The enlarged views are displayed in the insets. Cellular components are labelled accordingly. Nascent thylakoid: NT, chloroplast envelope: CE, chloroplast 70S ribosomes: 70S, cytoplasmic 80S ribosomes: 80S.

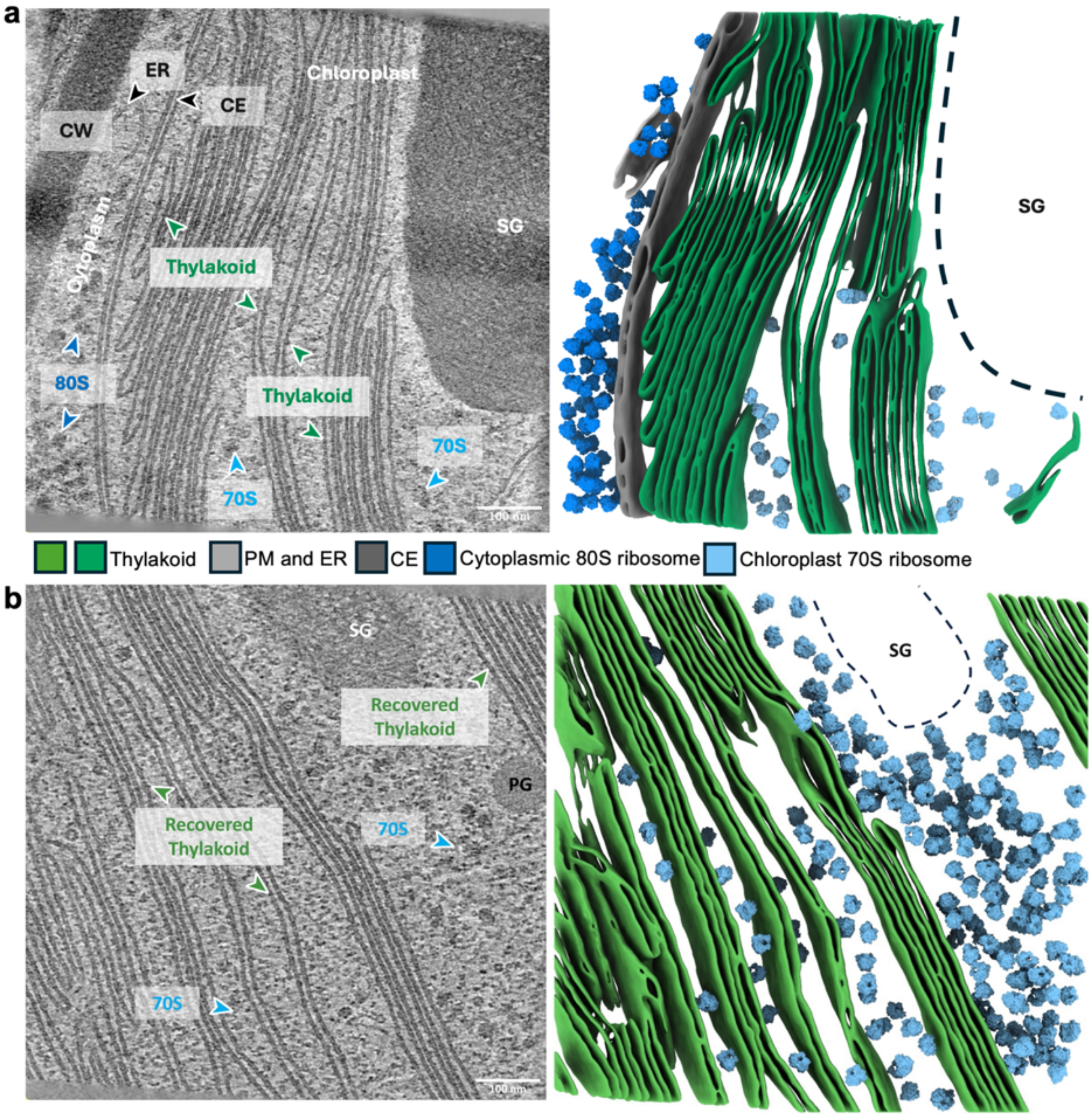

**Fig. S8. Cryo-ET of recovered Δ*chlL* and WT cells. a**, A representative tomographic slice and segmented volume of the thylakoid in WT. Cellular components are coloured and labelled accordingly. **b**, A representative tomographic slice and segmented volume of the thylakoid in Δ*chlL* after 24 h of light treatment. Nascent thylakoid: NT, chloroplast envelope: CE, plasma membrane: PM, cell wall: CW, chloroplast 70S ribosomes: 70S, cytoplasmic 80S ribosomes: 80S, endoplasmic reticulum: ER, plastoglobule: PG, and starch granule: SG.

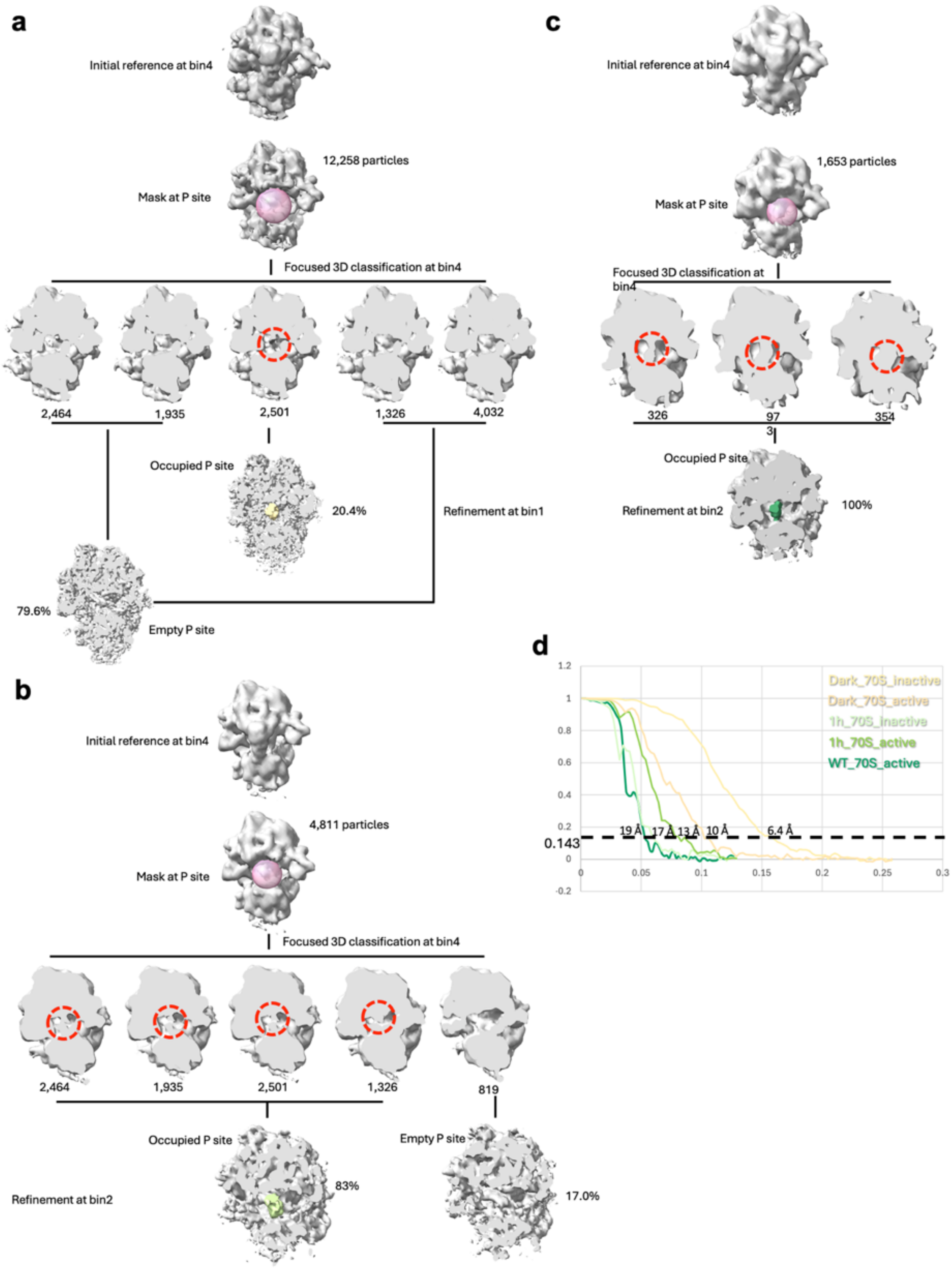

**Fig. S9. Workflow of subtomogram averaging of chloroplast 70S ribosomes in Δ*chlL* and WT. a**, Classification of chloroplast 70S ribosomes in dark-treated Δ*chlL* cells. **b**, Classification of chloroplast 70S ribosomes in 1-h light treated Δ*chlL* cells. **c**, Classification of chloroplast 70S ribosomes in dark-treated WT cells. The P site is masked (pink sphere) for focused classification and indicated by the red dotted circles after the classification. **d**, FSC plot of chloroplast 70S ribosomes, the resolution is indicated at gold-standard 0.143 FSC cut-off.

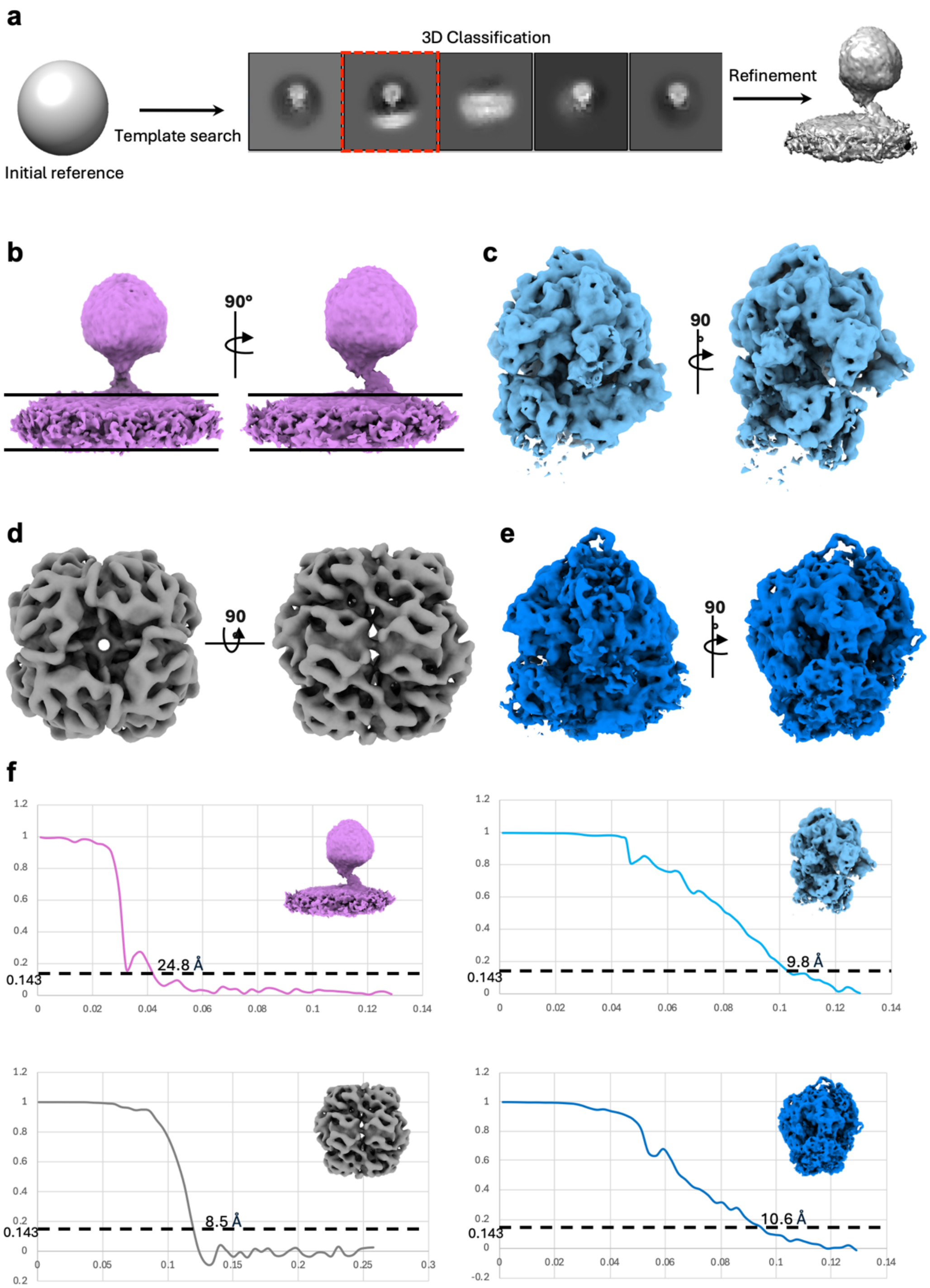

**Fig. S10. Subtomogram averaging of macromolecules in Δ*chlL*. a**, Workflow of bias-mitigated template matching for locating F-ATPase. A featureless sphere is employed as the initial template and the class with the stator and membrane densities is selected as true positive. **b**, Orthogonal views of in-cell structure of membrane-bound F-ATPase, the membrane is indicated by two parallel black lines. **c**, Orthogonal views of in-cell structure of chloroplast 70S ribosome. **d**, Orthogonal views of in-cell structure of pyrenoid RuBisCO. **e**, Orthogonal views of in-cell structure of cytoplasmic 80S ribosome. **f**, FSC plots of subtomogram averaged maps. The resolution is indicated at gold-standard 0.143 FSC cut-off.

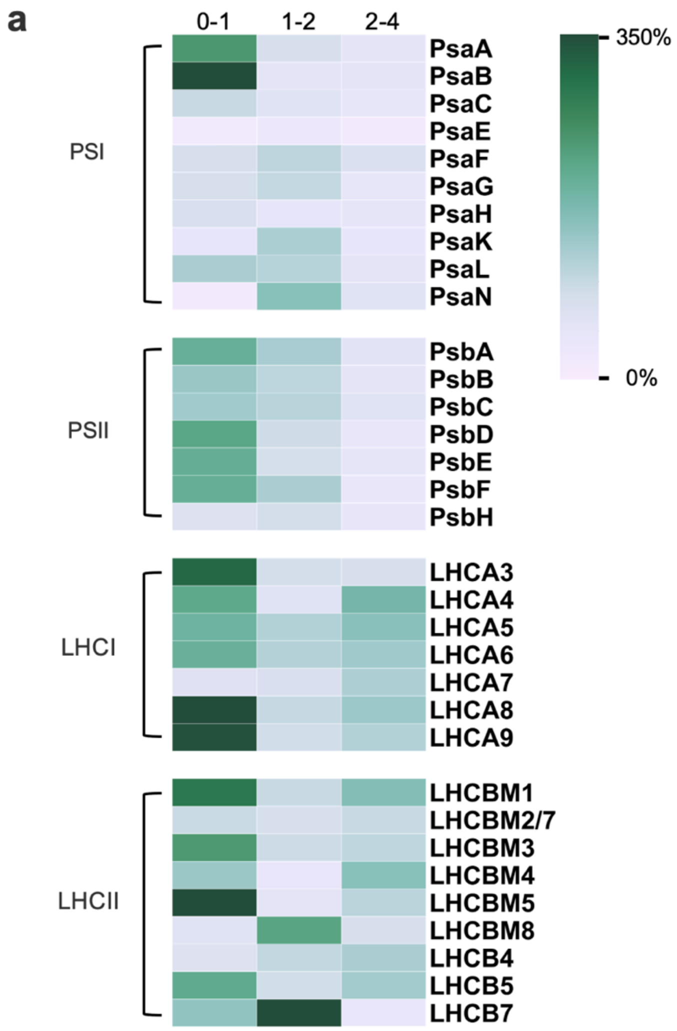
a
0-1
1-2
2-4
PSI
PsaA
PsaB
PsaC
PsaE
PsaF
PsaG
PsaH
PsaK
PsaL
PsaN
PSII
PsbA
PsbB
PsbC
PsbD
PsbE
PsbF
PsbH
LHCI
LHCA3
LHCA4
LHCA5
LHCA6
LHCA7
LHCA8
LHCA9
LHCII
LHCBM1
LHCBM2/7
LHCBM3
LHCBM4
LHCBM5
LHCBM8
LHCB4
LHCB5
LHCB7
350%
0%

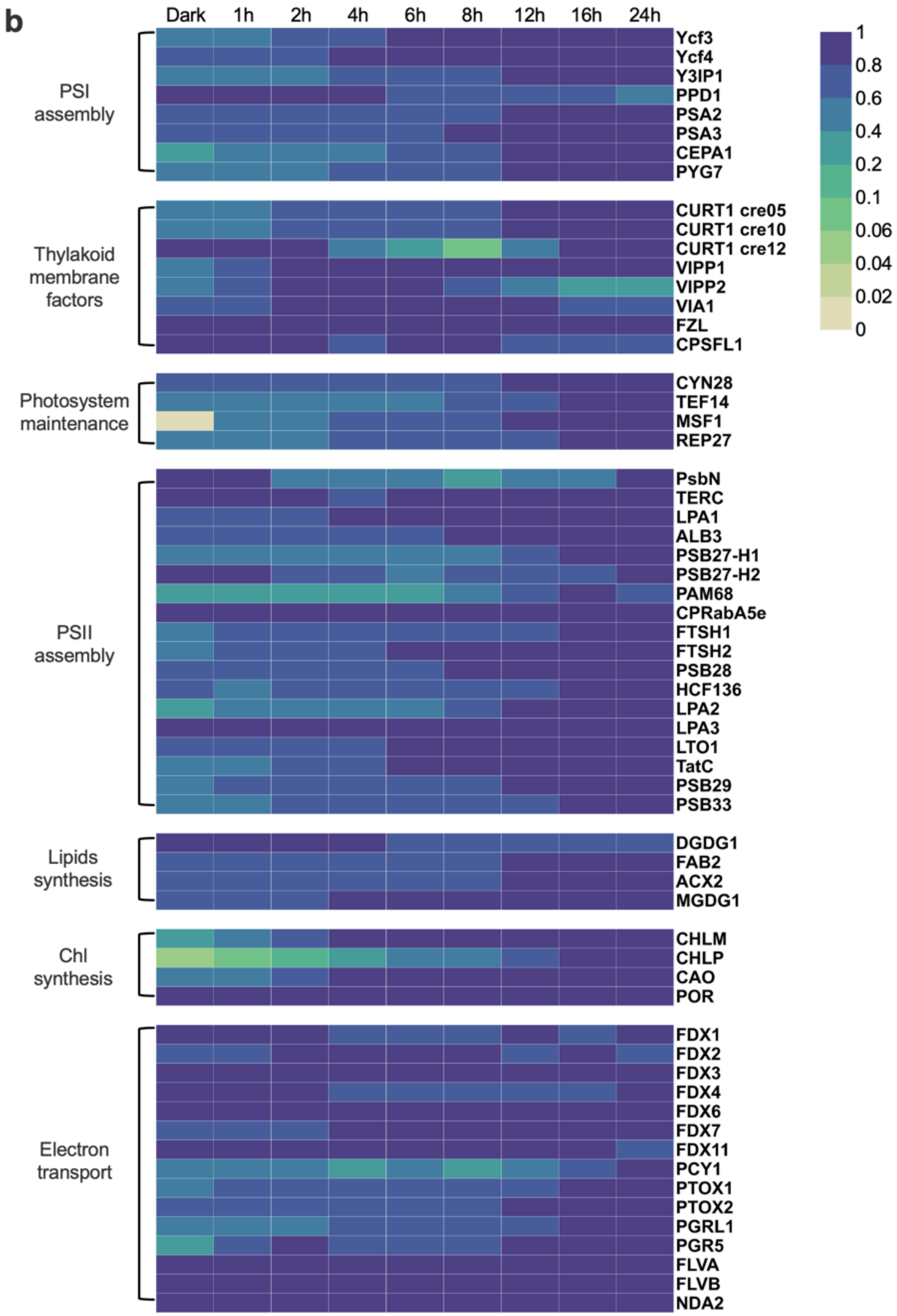
b
Dark
1h
2h
4h
6h
8h
12h
16h
24h
PSI assembly
Ycf3
Ycf4
Y3IP1
PPD1
PSA2
PSA3
CEPA1
PYG7
Thylakoid membrane factors
CURT1 cre05
CURT1 cre10
CURT1 cre12
VIPP1
VIPP2
VIA1
FZL
CPSFL1
Photosystem maintenance
CYN28
TEF14
MSF1
REP27
PSII assembly
PsbN
TERC
LPA1
ALB3
PSB27-H1
PSB27-H2
PAM68
CPRabA5e
FTSH1
FTSH2
PSB28
HCF136
LPA2
LPA3
LTO1
TatC
PSB29
PSB33
Lipids synthesis
DGDG1
FAB2
ACX2
MGDG1
Chl synthesis
CHLM
CHLP
CAO
POR
Electron transport
FDX1
FDX2
FDX3
FDX4
FDX6
FDX7
FDX11
PCY1
PTOX1
PTOX2
PGRL1
PGR5
FLVA
FLVB
NDA2
1
0.8
0.6
0.4
0.2
0.1
0.06
0.04
0.02
0

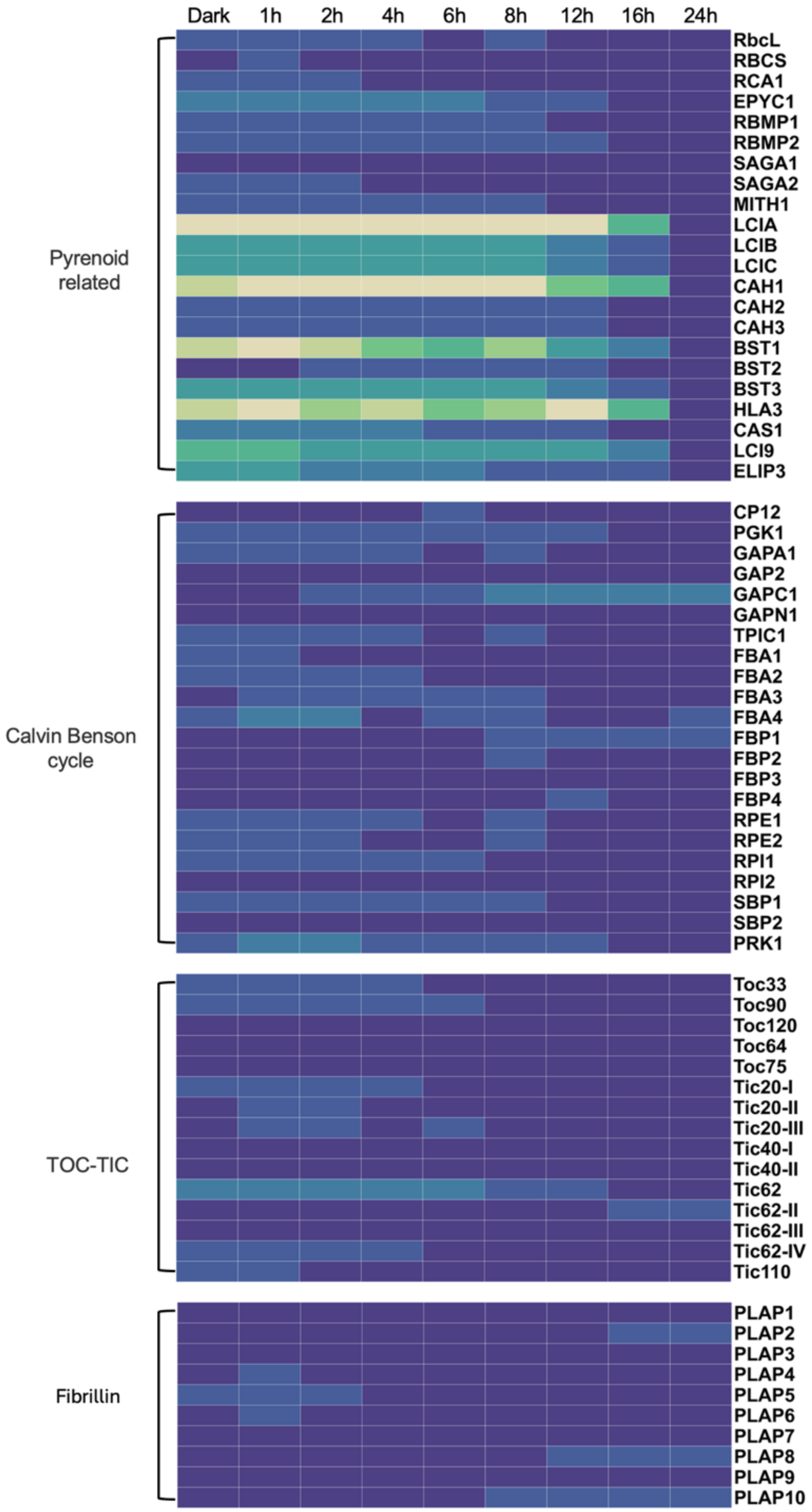
Dark
1h
2h
4h
6h
8h
12h
16h
24h
Pyrenoid related
RbcL
RBCS
RCA1
EPYC1
RBMP1
RBMP2
SAGA1
SAGA2
MITH1
LCIA
LCIB
LCIC
CAH1
CAH2
CAH3
BST1
BST2
BST3
HLA3
CAS1
LCI9
ELIP3
Calvin Benson cycle
CP12
PGK1
GAPA1
GAP2
GAPC1
GAPN1
TPIC1
FBA1
FBA2
FBA3
FBA4
FBP1
FBP2
FBP3
FBP4
RPE1
RPE2
RPI1
RPI2
SBP1
SBP2
PRK1
TOC-TIC
Toc33
Toc90
Toc120
Toc64
Toc75
Tic20-I
Tic20-II
Tic20-III
Tic40-I
Tic40-II
Tic62
Tic62-II
Tic62-III
Tic62-IV
Tic110
Fibrillin
PLAP1
PLAP2
PLAP3
PLAP4
PLAP5
PLAP6
PLAP7
PLAP8
PLAP9
PLAP10

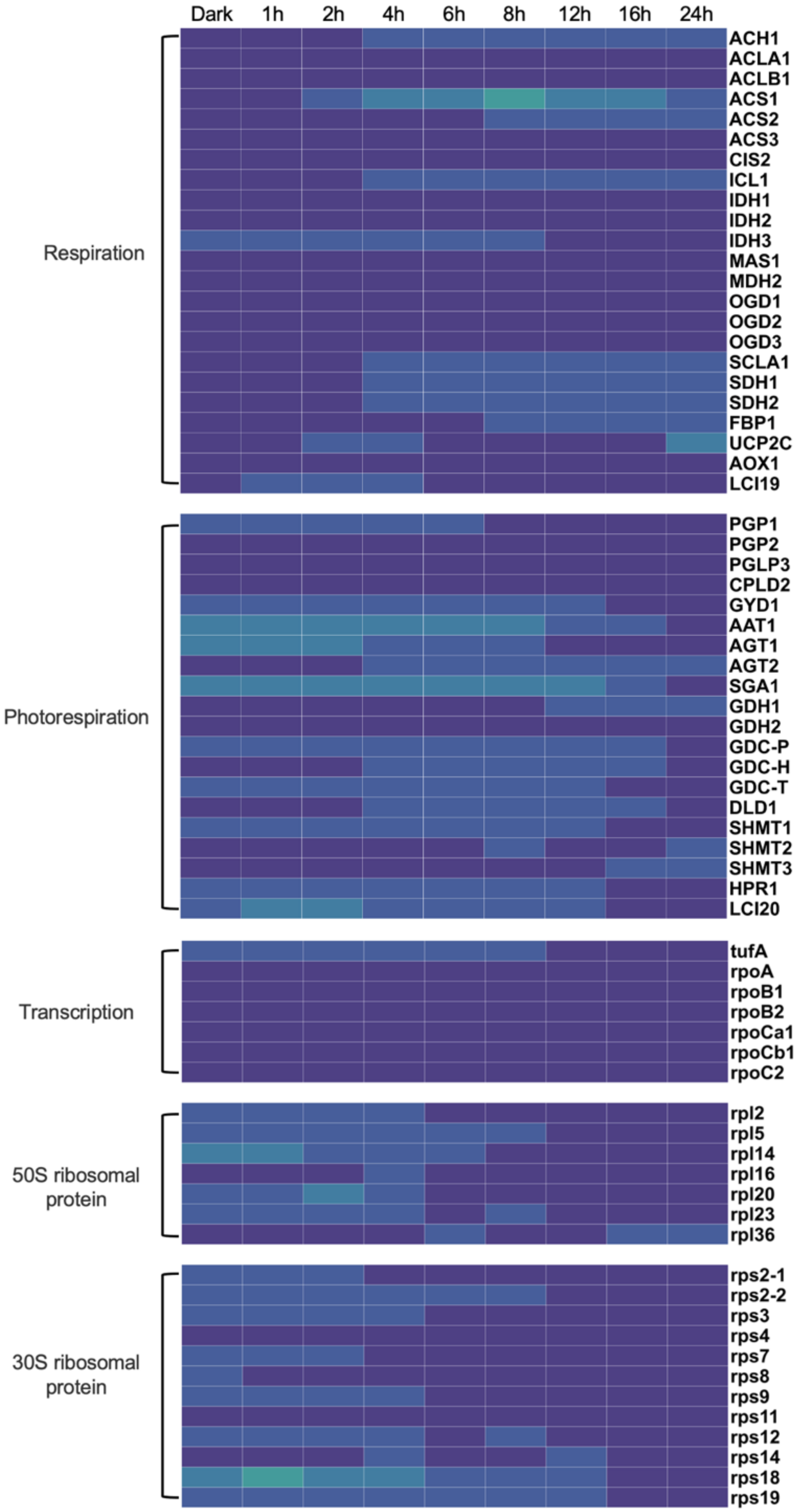
Dark
1h
2h
4h
6h
8h
12h
16h
24h
Respiration
ACH1
ACLA1
ACLB1
ACS1
ACS2
ACS3
CIS2
ICL1
IDH1
IDH2
IDH3
MAS1
MDH2
OGD1
OGD2
OGD3
SCLA1
SDH1
SDH2
FBP1
UCP2C
AOX1
LCI19
Photorespiration
PGP1
PGP2
PGLP3
CPLD2
GYD1
AAT1
AGT1
AGT2
SGA1
GDH1
GDH2
GDC-P
GDC-H
GDC-T
DLD1
SHMT1
SHMT2
SHMT3
HPR1
LCI20
Transcription
tufA
rpoA
rpoB1
rpoB2
rpoCa1
rpoCb1
rpoC2
50S ribosomal protein
rpl2
rpl5
rpl14
rpl16
rpl20
rpl23
rpl36
30S ribosomal protein
rps2-1
rps2-2
rps3
rps4
rps7
rps8
rps9
rps11
rps12
rps14
rps18
rps19

**Fig. S11. Changes in the protein abundance during thylakoid biogenesis. a**, The accumulation rates of individual proteins were calculated as the increment per hour to the previous sample, from 0-1h, 1-2h and 2-4h. Proteins resyntheses from relatively low levels were selected to highlight their earlier fold changes. **b**, Photosystem assembly factors, photosystem repair factors, membrane shaping factors, lipid synthases, Chl synthases, electron transport proteins, pyrenoid-related proteins, Calvin Benson cycle enzymes, TOC-TIC translocon system subunits, fibrillins, transcription, translation and proteins involved in respiration and photorespiration were manually grouped and normalized to the maximum level. Note that some homologous proteins have multiple names: Y3IP1/CGL59; PYG7/CGL71/Ycf37; PsbN/PBF1; Psb29/THF1; Psb33/TEF5/LIL8; LCI16/ELIP3; RBMP1/BST4; GDC-P/GCSP1; GDC-H/GCSH1; GDC-T/GCST1. CURT1 homologues in C. reinhardtii are located on chromosomes 5, 10, and 12, and were recently named CURT1A, CURT1C, and CURT1B.

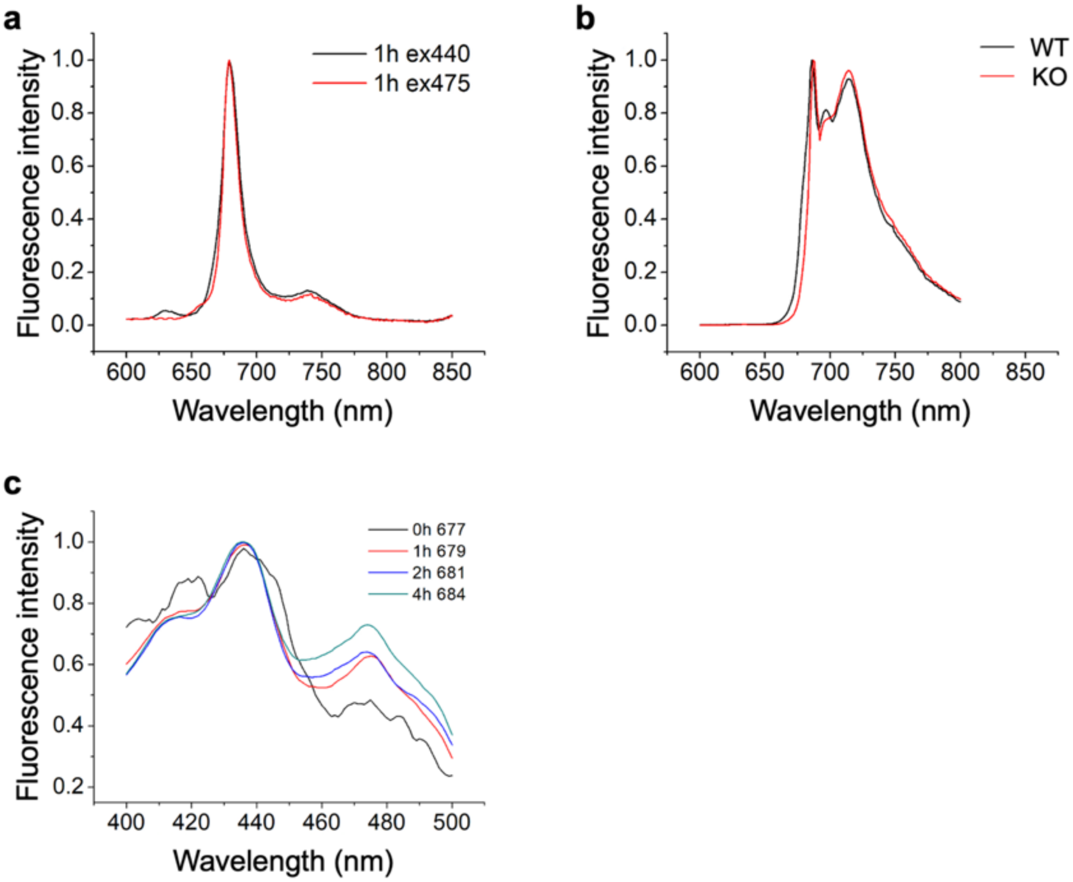

**Fig. S12. 77K fluorescence excitation and emission spectra. a**, Fluorescence emission spectra of Δ*chlL* samples treated with 1 h light, excited at 440 nm for Chl *a* and 475nm for Chl *b*, normalized to the maximum intensity. **b**, Fluorescence emission spectra of CC-1690 (WT) and Δ*chlL* (KO) grown under illumination, excited at 440 nm, normalized to the maximum intensity. **c**, Fluorescence excitation spectra of the main PSII peak of 0-4 h cells, normalized to the maximum intensity.

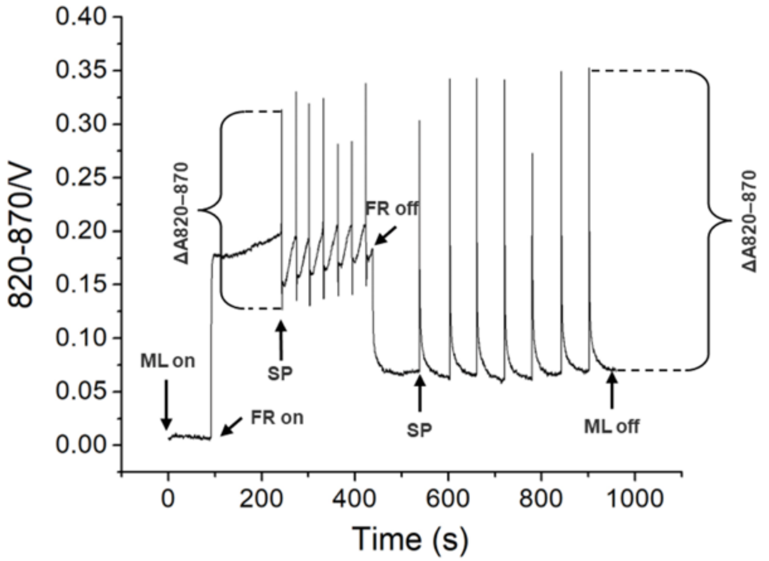

**Fig. S13. P700 slow kinetics measuring program.** Schematic diagram of the procedure for measuring slow kinetics of P700 using the PAM-KLAS system. ML, measure light, FR, far-red light, SP, saturation pulse.

**Table S1. Cryo-FIB lamella preparation and cryo-SEM.**

| Microscope | Conventional cryo-FIB/SEM Aquilos 2 | Plasma cryo-FIB/SEM Arctis | Plasma cryo-FIB/SEM Helios G4 Hydra |
|---|---|---|---|
| **Voltage (keV)** | 30 | 30 | 30 |
| **Ion beam source** | Gallium | Argon | Argon |
| **Sputtering coating prior to milling (seconds)** | No | 12 | 50 before and after GIS |
| **GIS coating time (second)** | 30 | 50 | 50 |
| **Bulk milling current** | N/A | N/A | N/A |
| **Milling current** | 0.1-0.5 nA | 0.74-2 nA | 2 nA and 200 pA for opening, 30 pA for milling |
| **Polishing current** | 30 pA | 60 pA | N/A |
| **Sputtering coating post polishing (seconds)** | No | No | No |
| **Fluorescence microscope** | METEOR (50 ×) | iFLM (100 × ) | METEOR (50 ×) |
| **Number of lamellae** | 46 | 39 | N/A, volume imaging |

**Table S2. Cryo-ET data collection.**

| Sample | Lamellae of WT | Lamellae of ∆*chlL* dark | Lamellae of ∆*chlL* 1h | Lamellae of ∆*chlL* 24h |
|---|---|---|---|---|
| **Microscope** | FEI Titan Krios G3 | FEI Titan Krios G3 | FEI Titan Krios G3 | FEI Titan Krios G3 |
| **Voltage (keV)** | 300 | 300 | 300 | 300 |
| **Detector** | Falcon 4i | Falcon 4i | Falcon 4i | Falcon 4i |
| **Energy-filter** | Selectris X | Selectris X | Selectris X | Selectris X |
| **Slit width (eV)** | 10 | 10 | 10 | 10 |
| **Super-resolution mode** | No | No | No | No |
| **Physical pixel size (Å/pixel)** | 1.978 | 1.94 | 1.94 | 1.9026 |
| **Defocus range (µm)** | -3 to -5, increment 0.3 | -3 to -5, increment 0.3 | -3 to -5, increment 0.3 | -3 to -5, increment 0.3 |
| **Acquisition scheme** | Dose-Symmetric, tilt span of 54º, 3º step, group 3 | Dose-Symmetric, tilt span of 54º, group 3 | Dose-Symmetric, tilt span of 54º, group 3 | Dose-Symmetric, tilt span of 54º, group 3 |
| **Total dose (electrons/Å$^2$)** | 129.5 | 129.5 | 129.5 | 129.5 |
| **Number of frames** | 10 | 10 | 10 | 10 |
| **Number of lamellae** | 39 | 17 | 21 | 8 |
| **Number of tomograms** | 118 | 152 | 163 | 31 |

**Table S3. Subtomogram averaging of macromolecules.**

| **Complexes** | **F-ATPase** | **Δ*chlL* (dark) inactive 70S ribosome** | **Δ*chlL* (dark) active 70S ribosome** | **Δ*chlL* (1h) inactive 70S ribosome** | **Δ*chlL* (1h) active 70S ribosome** | **WT (dark) active 70S ribosome** | **Cytoplasmic 80S ribosome** | **Pyrenoid RuBisCO** |
|---|---|---|---|---|---|---|---|---|
| **Particle number** | 2,026 | 9,757 | 2,501 | 819 | 3,992 | 1,653 | 2,950 | 21,560 |
| **Final resolution by gold-standard FSC cut (Å)** | 24.8 | 6.4 | 10.0 | 16.9 | 13.3 | 19.5 | 10.6 | 8.5 |

**Table S4. List of photosynthetic protein complex subunits.** Proteins identified by mass spectrometry in this study are in bold. This table is based on the *Chlamydomonas reinhardtii* chromosome whole genome shotgun sequence (NC_057004-NC_057020), The Chlamydomonas Genome Project v6 [29], chloroplast genome (NC_005353), Uniport and Chlamylibrary [30]. Note that PsbN under this nomenclature is not a structural component, PetO under this nomenclature was reported as a relatively independent component of cytochrome $b_6f$ but was not identified at high resolution, LHCBM2 and LHCBM7 are indistinguishable by mass spectrometry.

| Subunit | Locus tag | Cre ID | Protein accession | Description |
|---|---|---|---|---|
| **PsaA** | ChreCp019 | CreCp.g802280-CreCp.g802282 | P12154 | Photosystem I P700 chlorophyll *a* apoprotein A1 |
| **PsaB** | ChreCp048 | CreCp.g802312 | P09144 | Photosystem I P700 chlorophyll a apoprotein A2 |
| **PsaC** | ChreCp067 | CreCp.g802332 | Q00914 | Photosystem I iron-sulfur center |
| **PSAD** | CHLRE_05g238332v5 | Cre05.g238332 | Q5NKW4 | Photosystem I reaction center subunit II, chloroplastic |
| **PSAE** | CHLRE_10g420350v5 | Cre10.g420350 | A8ICV4 | Photosystem I reaction center subunit IV |
| **PSAF** | CHLRE_09g412100v5 | Cre09.g412100 | A8J4S1 | Photosystem I reaction center subunit III |
| **PSAG** | CHLRE_12g560950v5 | Cre12.g560950 | A0A2K3D5H4, P14224 | Photosystem I reaction center subunit V, chloroplastic |
| **PSAH** | CHLRE_07g330250v5 | Cre07.g330250 | A8IH77,P13352 | Photosystem I reaction center subunit VI, chloroplastic |
| **PSAI** | CHLRE_03g165100v5 | Cre03.g165100 | A8IFG7 | Photosystem I reaction center subunit VIII |
| **PsaJ** | ChreCp061 | CreCp.g802326 | P59777 | Photosystem I reaction center subunit IX |
| **PSAK** | CHLRE_17g724300v5 | Cre17.g724300 | P14225 | Photosystem I reaction center subunit psaK, chloroplastic |
| **PSAL** | CHLRE_12g486300v5 | Cre12.g486300 | A8IL32 | Photosystem I reaction center subunit L |
| **PSAN** | CHLRE_02g082500v5 | Cre02.g082500 | Q9AXJ2 | Photosystem I reaction center subunit N, chloroplastic |
| **PSAO** | CHLRE_07g334550v5 | Cre07.g334550 | A8JCL6 | Photosystem I subunit O |

| | | | | |
|---|---|---|---|---|
| **PsbA** | ChreCp021, ChreCp057 | CreCp.g802285, CreCp.g802321 | P07753 | Photosystem II protein D1 |
| **PsbB** | ChreCp032 | CreCp.g802297 | P37255 | Photosystem II CP47 reaction center protein |
| **PsbC** | ChreCp066 | CreCp.g802331 | P10898 | Photosystem II CP43 reaction center protein |
| **PsbD** | ChreCp064 | CreCp.g802329 | P06007 | Photosystem II D2 protein |
| **PsbE** | ChreCp040 | CreCp.g802304 | P48268 | Cytochrome b559 subunit alpha |
| **PsbF** | ChreCp043 | CreCp.g802307 | Q08363 | Cytochrome b559 subunit beta |
| **PsbH** | ChreCp029 | CreCp.g802294 | P22666 | Photosystem II reaction center protein H |
| **PsbI** | ChreCp051 | CreCp.g802315 | P59763 | Photosystem II reaction center protein I |
| **PsbJ** | ChreCp063 | CreCp.g802328 | O19930 | Photosystem II reaction center protein J |
| **PsbK** | ChreCp004 | CreCp.g802266 | A0A218N8C5, P18263 | Photosystem II reaction center protein K |
| **PsbL** | ChreCp044 | CreCp.g802308 | P32974 | Photosystem II reaction center protein L |
| **PsbM** | ChreCp026 | CreCp.g802290 | A0A218N984, P92277 | Photosystem II reaction center protein M |
| **PsbN** | ChreCp030 | CreCp.g802295 | Q06480 | Photosystem Biogenesis Factor 1 (PBF1) |
| **PSBO** | CHLRE_09g396213v5 | Cre09.g396213 | A8J0E4 | Oxygen-evolving enhancer protein 1 of photosystem II |
| **PSBP** | CHLRE_12g550850v5 | Cre12.g509050 | A0A2K3D661 | Oxygen-evolving enhancer protein 2 of photosystem II |
| **PSBQ** | CHLRE_08g372450v5 | Cre08.g372450 | A8JEV1 | Oxygen-evolving enhancer protein 3 of photosystem II |
| **PSBR** | CHLRE_06g261000v5 | Cre06.g261000 | A0A2K3DMP5 | Photosystem II protein PSBR, chloroplastic |
| **PSBS** | CHLRE_01g016750v5 | Cre01.g016600 | A8HPM5 | Photosystem II protein PSBS2 |
| **PsbT** | ChreCp031 | CreCp.g802296 | A0A218N8W1 | Photosystem II reaction center protein T |
| **PSBW** | CHLRE_11g801318v5 | Cre11.g801318 | Q9SPI9 | Photosystem II reaction center W protein, chloroplastic |

| | | | | |
|---|---|---|---|---|
| **PSBX** | CHLRE_02g082750v5, CHLRE_02g082852v5 | Cre02.g082750 | A8I846 | Photosystem II reaction center protein X |
| **PsbZ** | ChreCp027 | CreCp.g802291 | A0A218N8D7, P92276 | Photosystem II reaction center protein Z, ycf9 |
| **Psb30** | ChreCp022 | CreCp.g802286 | P50370 | Photosystem II reaction center protein Psb30, ycf12 |
| **PetA** | ChreCp001 | CreCp.g802263 | P23577 | Cytochrome f |
| **PetB** | ChreCp008 | CreCp.g802270 | Q00471 | Cytochrome b6 |
| **PETC** | CHLRE_11g467689v5 | Cre11.g467689 | A8J9Y1 | Chloroplast cytochrome b6f Rieske iron-sulfur center subunit |
| **PetD** | ChreCp002 | CreCp.g802264 | P23230 | Cytochrome b6-f complex subunit 4 |
| **PetG** | ChreCp045 | CreCp.g802309 | Q08362 | Cytochrome b6-f complex subunit 5 |
| **PETM** | CHLRE_12g546150v5 | Cre12.g546150 | Q42496, A8IY88 | Cytochrome b6-f complex subunit 7, chloroplastic |
| **PetL** | ChreCp068 | CreCp.g802333 | A0A218N8J1 | Cytochrome b6-f complex subunit 6 |
| **PETO** | CHLRE_12g558900v5 | Cre12.g558900 | Q9LLC6, A8JGW2 | Cytochrome b6-f complex subunit petO, chloroplastic |
| **PETN** | CHLRE_16g650100v5 | Cre16.g650100 | P0C1D4 | Subunit of the chloroplast cytochrome b6f complex |
| **AtpA** | ChreCp050 | CreCp.g802314 | P26526 | ATP synthase subunit alpha, chloroplastic |
| **AtpB** | ChreCp058 | CreCp.g802323 | P06541 | ATP synthase subunit beta, chloroplastic |
| **ATPC** | CHLRE_06g259900v5 | Cre06.g259900 | A8HXL8, P12113 | ATP synthase gamma chain, chloroplastic |
| **ATPD** | CHLRE_11g467569v5 | Cre11.g467569 | A8JF15, Q42687 | ATP synthase delta chain, chloroplastic |
| **AtpE** | ChreCp023 | CreCp.g802287 | P07891 | ATP synthase epsilon chain, chloroplastic |
| **AtpF** | ChreCp054 | CreCp.g802318 | Q8HTL5 | ATP synthase subunit b, chloroplastic |

| | | | | |
|---|---|---|---|---|
| **ATPG** | CHLRE_11g481450 v5 | Cre11.g481450 | A0A2K3D8S8, A8J785 | ATP synthase subunit b', chloroplastic |
| **AtpH** | ChreCp053 | CreCp.g802317 | Q37304 | ATP synthase subunit c, chloroplastic |
| **AtpI** | ChreCp062 | CreCp.g802327 | O63075 | ATP synthase subunit a, chloroplastic |
| **LHCA1** | CHLRE_06g283050 v5 | Cre06.g283050 | Q05093 | Chlorophyll a-b binding protein, chloroplastic |
| **LHCA2** | CHLRE_12g508750 v5 | Cre12.g508750 | A8IKC8 | Chlorophyll a-b binding protein, chloroplastic |
| **LHCA3** | CHLRE_11g467573 v5 | Cre11.g467573 | Q75VY9 | Chlorophyll a-b binding protein, chloroplastic |
| **LHCA4** | CHLRE_10g452050 v5 | Cre10.g452050 | Q75VZ0 | Chlorophyll a-b binding protein, chloroplastic |
| **LHCA5** | CHLRE_10g425900 v5 | Cre10.g425900 | Q75VY8 | Chlorophyll a-b binding protein, chloroplastic |
| **LHCA6** | CHLRE_06g278213 v5 | Cre06.g278213 | Q75VY6 | Chlorophyll a-b binding protein, chloroplastic |
| **LHCA7** | CHLRE_16g687900 v5 | Cre16.g687900 | Q84Y02 | Chlorophyll a-b binding protein, chloroplastic |
| **LHCA8** | CHLRE_06g272650 v5 | Cre06.g272650 | Q75VY7 | Chlorophyll a-b binding protein, chloroplastic |
| **LHCA9** | CHLRE_07g344950 v5 | Cre07.g344950 | A8ITV3 | Chlorophyll a-b binding protein, chloroplastic |
| **LHCBM1** | CHLRE_01g066917 v5 | Cre01.g066917 | Q93VE0 | Chlorophyll a-b binding protein, chloroplastic |
| **LHCBM2** | CHLRE_12g548400 v5 | Cre12.g548400 | Q93WE0 | Chlorophyll a-b binding protein, chloroplastic |
| **LHCBM3** | CHLRE_04g232104 v5 | Cre04.g232104 | Q93WL4 | Chlorophyll a-b binding protein, chloroplastic |
| **LHCBM4** | CHLRE_06g283950 v5 | Cre06.g283950 | A8J264 | Chlorophyll a-b binding protein, chloroplastic |
| **LHCBM5** | CHLRE_03g156900 v5 | Cre03.g156900 | Q9ZSJ4 | Chlorophyll a-b binding protein, chloroplastic |
| **LHCBM6** | CHLRE_06g285250 v5 | Cre06.g285250 | A8J287 | Chlorophyll a-b binding protein, chloroplastic |
| **LHCBM7** | CHLRE_12g548950 v5 | Cre12.g548950 | Q9AXF6 | Chlorophyll a-b binding protein, chloroplastic |
| **LHCBM8** | CHLRE_06g284250 v5 | Cre06.g284250 | A8J270 | Chlorophyll a-b binding protein, chloroplastic |
| **LHCBM9** | CHLRE_06g284200 v5 | Cre06.g284200 | Q8S3T9 | Chlorophyll a-b binding protein, chloroplastic |

| | | | | |
|---|---|---|---|---|
| **LHCBM10** | Not reported | | Q8S3U0 | Chlorophyll a-b binding protein, chloroplastic |
| **CP29, LHCB4** | CHLRE_17g720250v5 | Cre17.g720250 | Q93WD2 | Chlorophyll a-b binding protein, chloroplastic |
| **CP26, LHCB5** | CHLRE_16g673650v5 | Cre16.g673650 | Q9FEK6 | Chlorophyll a-b binding protein, chloroplastic |
| **LHCB7, LHCQ** | CHLRE_02g110750v5 | Cre02.g110750 | A0A2K3E2Y8 | Chlorophyll a-b binding protein, chloroplastic |
